# An organoid platform reveals MEK-PARP co-targeting to enhance radiation response in rectal cancer

**DOI:** 10.1101/2024.06.06.597640

**Authors:** Qiyun Xiao, Julian E. Riedesser, Theresa Mulholland, Zhenchong Li, Jonas Buchloh, Philipp Albrecht, Moying Li, Nachiyappan Venkatachalam, Olga Skabkina, Anna Klupsch, Ella Eichhorn, Li Wang, Sebastian Belle, Nadine Schulte, Daniel Schmitz, Matthias F. Froelich, Erica Valentini, Kim E. Boonekamp, Yvonne Petersen, Thilo Miersch, Elke Burgermeister, Carsten Herskind, Marlon R. Veldwijk, Christoph Brochhausen, Robert Ihnatko, Jeroen Krijgsveld, Ina Kurth, Michael Boutros, Matthias P. Ebert, Tianzuo Zhan, Johannes Betge

**Author notes:** Equal contribution (Qiyun Xiao, Julian E. Riedesser, Theresa Mulholland, Tianzuo Zhan, Johannes Betge). TZ and JB jointly directed this work. Correspondence to TZ and JB.

## Abstract

Locally advanced rectal cancer is usually treated by neoadjuvant chemoradiotherapy. However, tumor response rates to this treatment vary greatly. Thus, most patients do not reach a complete remission and have to undergo tumor resection. In the present study, we introduce a patient-derived rectal cancer organoid platform that reflects clinical radiosensitivity and use this to screen 1596 drug-radiation combinations. We identify inhibitors of RAS-MAPK signaling, especially MEK inhibitors, strongly synergizing with radiation response. Mechanistically, MEK inhibitors suppressed radiation-induced activation of RAS-MAPK signaling, and selectively downregulated the homologous recombination DNA repair pathway component RAD51, thereby achieving radio-enhancement. Through testing drug-drug-radiation combinations in organoids and cell lines, we identified synergism between PARP and MEK inhibitors to further enhance the effect of radiation. Our data support clinical testing of combined MEK and PARP inhibition with radiotherapy in locally advanced rectal cancers.

**Graphical Abstract:** 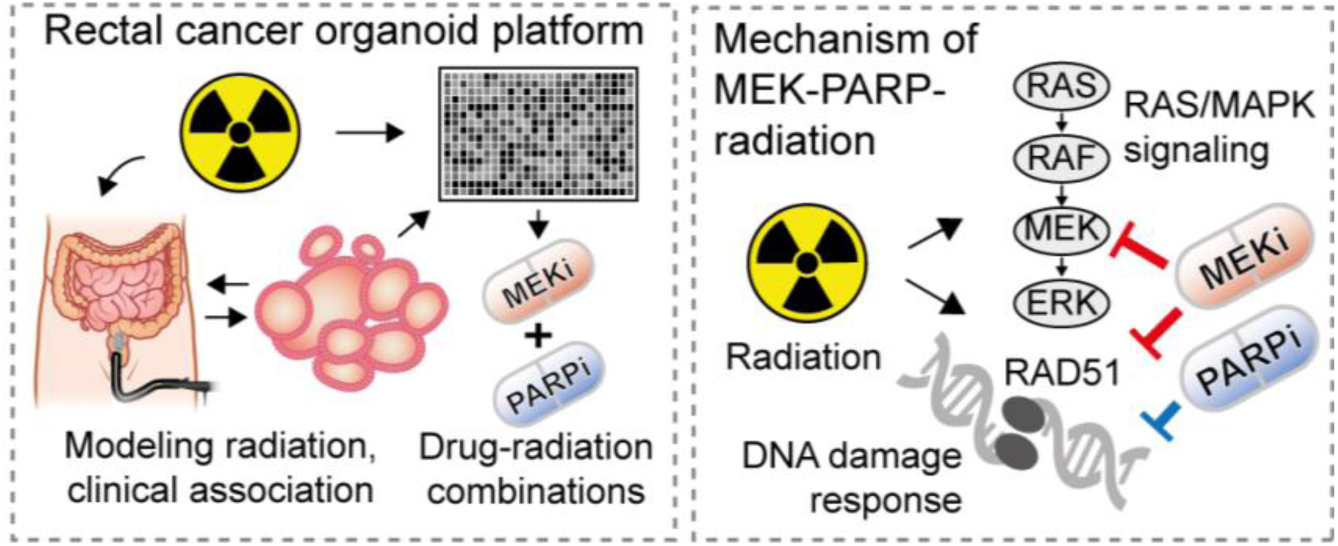

---

Colorectal cancer (CRC) stands as a leading cause of cancer-related mortality.^1^ Over one third of CRC originates in the rectum, often presenting at a locally advanced stage, which is defined as Union for International Cancer Control (UICC) classification T3/T4 (invasion beyond muscular layers) and/or node positive disease. The current standard of care for most locally advanced rectal cancers is neoadjuvant chemoradiotherapy, followed by surgical resection of the tumor.^2^ Introduction of neoadjuvant chemoradiotherapy led to improved local tumor control rates in clinical trials.^3^ The most commonly applied regimens include a combination of long-course radiation with intravenous or oral fluoropyrimidine and, more recently, the addition of consolidation chemotherapy with 5-fluorouracil and oxaliplatin, termed total neoadjuvant therapy (TNT).^4^ The response to neoadjuvant chemoradiotherapy varies significantly between individuals, ranging from complete responses without detectable tumor residues to non-response. Patients who achieve complete response have an improved overall survival and may avoid surgical resection of the rectum, which can be severely debilitating. While standard neoadjuvant chemoradiotherapy has resulted in complete response rates around 15%,^5^ intensified regimens such as TNT can significantly enhance response rates. However, this success is achieved at the expense of increased toxicity, which is caused by the broad mode-of-action of conventional chemotherapeutic agents, notably neurotoxicity induced by oxaliplatin.^6^ Tailored approaches that target tumor-specific alterations to enhance radiosensitivity have not been introduced into clinical practice, despite promising preclinical results with different small molecule drugs and antibodies.^7^ One of the underlying reasons is the absence of suitable tumor models that adequately reflect the biological characteristics of rectal cancers, which are dominated by particularly high frequencies of RAS- and WNT pathway mutations.^8^ Recently, patient-derived organoids have been introduced as models that can recapitulate the tumor biology of many cancer types and their response to different therapeutic modalities.^9^ In particular, studies have shown associations of rectal cancer organoids’ response to radiation with response of corresponding tumors.^10–12^ So far, however, rectal cancer organoid platforms have not been exploited to systematically screen for novel drug candidates that can enhance the response of rectal cancers to radiotherapy.

In this study, we establish a rectal cancer organoid platform that recapitulates clinical radiosensitivity and use it to perform large-scale drug screens to identify drugs synergizing with radiation therapy. We observed that inhibitors of the RAS-MAPK pathway, in particular MEK inhibitors, can strongly increase sensitivity of rectal cancer organoids and CRC cell lines to radiation. Mechanistically, we find that radiotherapy induces an activation of RAS-MAPK signaling which could be suppressed by MEK inhibition. Moreover, MEK inhibitors downregulate RAD51 protein levels, a key component of the DNA repair machinery. Accordingly, we find that MEK inhibitors synergize with PARP1/2 inhibitors in reducing tumor cell viability, and the combination of these two agents can further enhance the effectiveness of radiotherapy in CRC cell lines and organoids.

## Results

### An organoid platform recapitulates essential aspects of rectal cancer

To model cancer biology and identify novel treatment options for rectal cancer, we established an organoid-based platform and living biobank. Organoids were generated from pre-treatment endoscopic biopsies from patients with rectal cancers of different UICC/TNM stages (Fig. 1A, Table S1). They showed heterogeneous morphologies and molecular alterations that are characteristic for CRC (Fig. 1B).^13^ Previous studies have demonstrated associations between clinical signs of radiation response in rectal cancer patients and radiation response of corresponding rectal cancer organoids, using various protocols.^10–12^ We established a short-term, standardized, robot-assisted radiation protocol for organoids, based on our previously published high-throughput screening platform.^14^ By exposing tumor organoid cultures to different doses of radiation, we observed a clear dose-dependency of organoid viability and a high degree of variation in radiosensitivity between different patient-donors (Fig. 1C-E, Fig. S1). The variation in response was not explained by frequent mutations found in the organoids, consistent with previous findings (Fig S1).^8^ Clinical response to chemoradiotherapy determined by magnetic resonance tomography (MRI) regression grading showed a significant association (almost perfect except for one outlier) with the response of corresponding patient-derived rectal cancer organoids to radiation (Fig. 1D-F). Additionally, analysis of regression grade in post-radiation tumor resection specimens, as well as analysis of changes in tumor length in MRI also showed a strong association (although not statistically significant) with organoid response (Fig. 1D-F, Fig. S1). Finally, endoscopic assessment of therapy response corresponded to the radiosensitivity of organoids in our assay (representative clinical images shown in Fig. 1F). These findings are in concordance with observations of previous studies and support the clinical and functional relevance of our organoid radiation assay in modeling radiosensitivity.

**Fig. 1:**
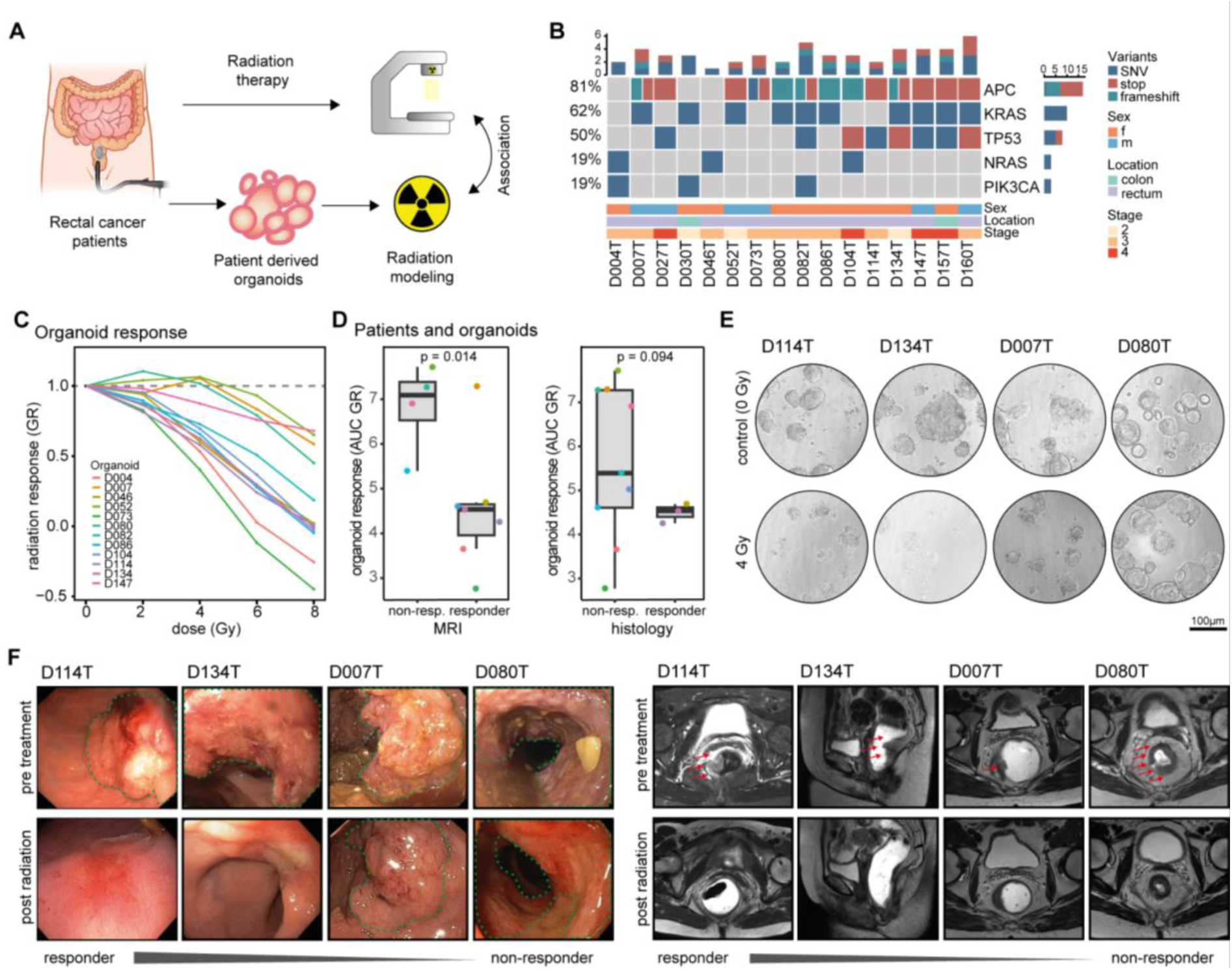
An organoid platform recapitulates essential aspects of rectal cancer. **A,** Schematic illustration of the rectal cancer organoid platform and approach for correlation of patient and organoid response. **B**, Driver mutations identified in patient-derived organoids. **C,** Radiation response of rectal cancer organoids to increasing doses of radiation. **D**, Analysis of organoid response to radiation according to donor patients’ rectal cancer response to radiation therapy assessed by MRI based regression grading (left, 11 evaluable cases, two-tailed Student’s t-test) and histopathological examination (right, 12 evaluable cases, two-tailed Welch’s t-test). **E**, Representative brightfield images of rectal cancer organoid cultures undergoing radiation. **F**, Representative endoscopy (left panel) and MRI (right panel) images from selected patients pre-treatment and post radiation therapy, sorted according to organoid response to radiation therapy.

### High-throughput screening in rectal cancer organoids identifies MAPK-signaling inhibitors as enhancers of radiation response

Radiation is usually combined with fluoropyrimidines as chemotherapeutic agents in the treatment of locally advanced rectal cancer. While fluoropyrimidines have been reported to exhibit radiosensitizing properties,^15^ their modes of action are broad and non-selective. Hence, novel drugs with stronger radiosensitizing effects are needed to achieve deeper responses and may spare patients from debilitating rectal resection. To screen for novel synergistic treatment combinations of radiation with medical therapies, we used our rectal cancer organoid biobank and further developed our semi-automated radiation workflow towards a high-throughput assay for combined drug and radiation therapy screening. Within this workflow, organoids underwent radiation and drug treatments in 384-well format in a short-term viability assay over nine days (Fig. 2A). We used ΔAUCs (area under the dose-response curve of non-irradiated - irradiated conditions) as a simple metric to screen for drugs that were enhanced by the radiation effect. Normalizing perturbations to the plate-specific dimethyl sulfoxide (DMSO) controls (i.e. separately for irradiated or non-irradiated plates) thereby revealed radio-enhancing effects, avoiding bias between the screened plates. We first used two organoid lines that were resistant to radiation therapy (D080T, D007T, compare Fig. 1) and applied radiation treatment in combination with a library of 224 drugs, mostly kinase inhibitors, in four concentrations (Fig. 2B, Fig. S2). In both organoid lines, we identified several kinase inhibitors that enhanced radiation effects and also compounds that diminished tumor cell killing upon radiation treatment (Fig. 2C-F). The enhancing effects were generally stronger in organoid line D080T than in D007T (Fig. 2C and E). Many compounds that conferred high radiation-induced killing in both organoid lines belonged to the class of MAPK signaling inhibitors (including inhibitors of EGFR, MEK or RAF, Fig. 2D and F). To validate these findings and to identify additional clinically available drugs enhancing radiation with the potential of a fast clinical translation, we performed further screening experiments. We used a library containing 140 cancer drugs, mostly FDA-approved, that could be meaningfully modeled in organoid experiments (i.e. including drugs with a mechanism directly targeting cancer cells, Fig. 2G). This library included every drug in five concentrations and was tested in 10 different rectal cancer organoid models (Fig. 2H-L). Most drugs showed similar responses in irradiated and non-irradiated conditions, while mainly inhibitors of the RAS-MAPK signaling pathway recurrently demonstrated enhanced killing together with radiation (Fig. 2H). Ranking drugs according to ΔAUC demonstrated that RAS-MAPK (EGFR, MEK) pathway inhibitors, as well as PARP inhibitors, which are known radiosensitizers in different tumors,^16^ were among the strongest hits (Fig. 2I-K, Fig. S3). The degree of radiosensitization varied between the different organoid models, likely representing the molecular heterogeneity of rectal cancers (Fig. 2L, Fig. S3). With respect to MEK inhibitors, seven out of ten organoid lines showed strongly enhanced tumor organoid killing in varying degrees, when combined with radiation (Fig. 2L). EGFR inhibitors exhibited similar, but not congruent profiles of radio-enhancement across different organoid lines compared to MEK inhibitors, while the profiles were distinct for PARP inhibitors (Fig. S3). We then tested associations of common molecular alterations in organoid lines with the level of radiosensitization. We found that organoid lines with wild-type *TP53* status generally showed drug responses that were more strongly modifiable by radiation, especially with MEK inhibitors, while no association of radiation-enhancement was observed with RAS mutation status (Fig. S3D-E). In conclusion, high-throughput drug-radiation combination screens with both kinase and clinical libraries independently showed a strong positive interaction of radiation with inhibitors of RAS-MAPK signaling, especially MEK inhibitors.

**Fig. 2:**
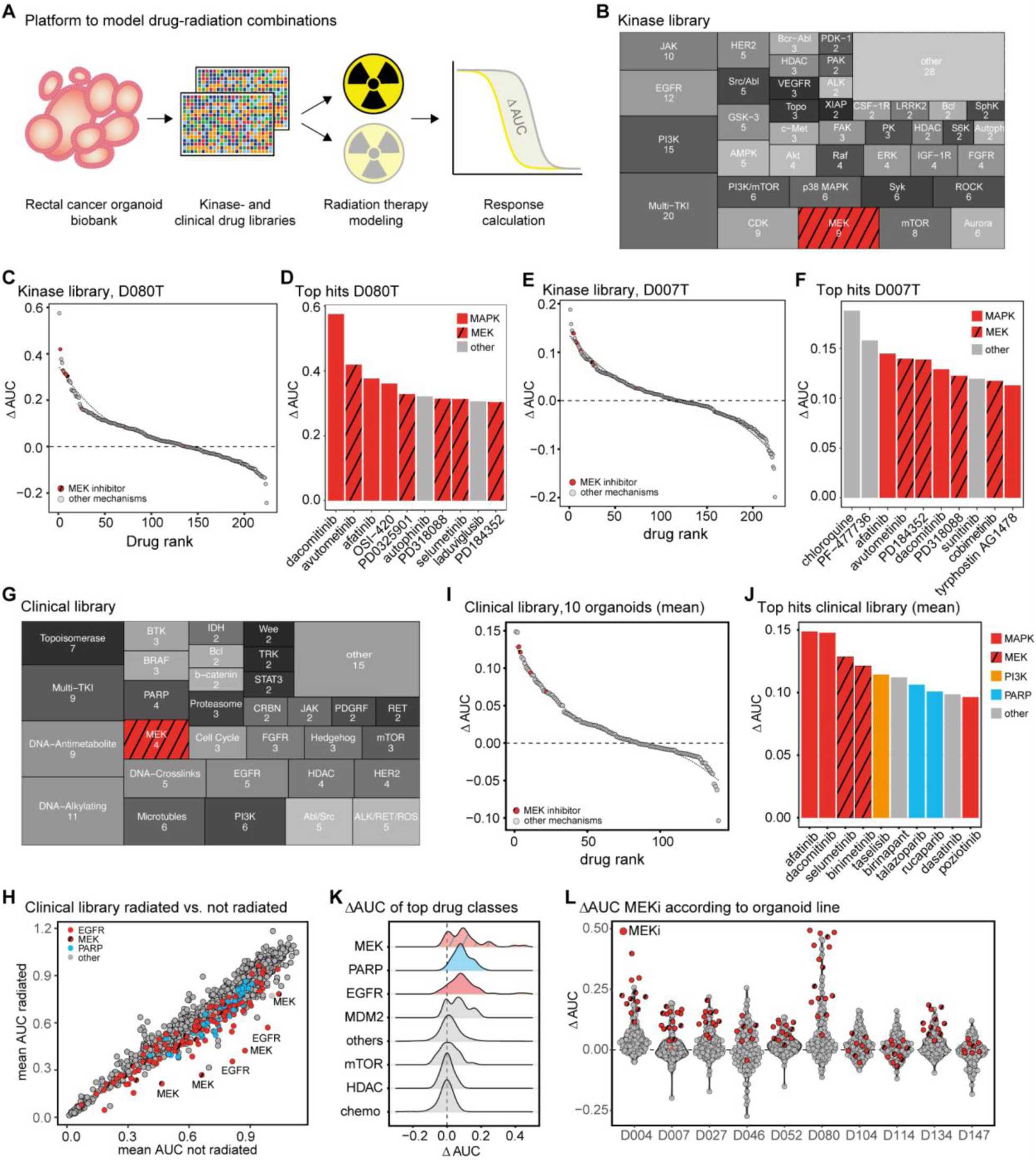
Drug screening identifies RAS-MAPK inhibitors to synergistically enhance radiation in rectal cancer models. **A,** Schematic representation of the drug-radiation screening workflow. Rectal cancer organoids were seeded in 384-well plates, drug perturbations with 4-5 concentrations and radiation (2 - 4 Gy) were performed on day 3, before viability was measured on day 9 after seeding. Interactions of drugs and radiation were analyzed by calculating the difference between areas under the dose-response-curves (ΔAUC values) between irradiated and non-irradiated conditions, each normalized to respective irradiated and non-irradiated DMSO controls on the same plates. **B,** Composition of the kinase drug library with 224 drugs, tested in 4 concentrations in two organoid lines. **C,** Ranking of differential effects of kinase inhibitors with or without radiation in tumor organoid D080T. **D,** Top 10 hits with highest radiation enhancement (ΔAUC values) in the kinase library screen of organoid line D080T. **E,** Ranking of differential effects of kinase inhibitors with or without radiation in cancer organoid D007T. **F,** Top 10 hits with strongest radiation enhancement (ΔAUC values) in the kinase library screen of organoid line D007T. **G,** Composition of the clinical cancer library consisting of 140 drugs. The drugs were administered in 5 concentrations and 10 organoid lines were tested. **H**, Mean area under the curve of all drugs tested in the clinical library in irradiated vs. non-irradiated conditions. **I**, Ranking the mean differential effects of clinical cancer drugs with or without radiation in 10 rectal cancer organoids. **J**, Top 10 hits with strongest radiation enhancement (mean ΔAUC values) in the clinical library screen with ten rectal cancer organoids. **K**, Distribution of ΔAUCs of selected groups of inhibitors. **L**, ΔAUCs of individual organoid lines, ΔAUCs of MEK inhibitors are highlighted.

### MEK inhibition is synergistic with radiation in CRC cell lines and organoids

Amongst the identified drug candidates, MEK-inhibitors (MEKi) showed the strongest enhancement by radiation. MEKi targets RAS-MAPK signaling downstream of oncogenic RAS mutations, which are highly prevalent in rectal cancers.^8^ In two previously tested organoid lines (D080T and D007T), radiation with 4 Gy combined with the FDA-approved MEKi trametinib resulted in a significant decrease of cell viability, which was significantly stronger in combination with radiation treatment (Fig. 3A-B). To prove a synergistic effect of MEKi with radiation, we calculated the expected combination response of both perturbations for each tested concentration using a Bliss independence model, as recently reported for drug-drug combinations.^17^ This revealed a clear excess of the experimentally observed combination response over the expected response, proving synergy between radiation and MEKi. Microscopy images of organoids showed corresponding phenotypes, with a reduction of organoid size and number after combination therapy (Fig. 3A-B). Replicative cell death is a major cause for the antineoplastic effect of radiotherapy. To allow for testing radiation effects over several cycles of cell proliferation, we performed complementary experiments in CRC cell lines using viability assays and gold-standard colony forming assays. We selected three commonly used CRC cell lines SW480, DLD1 and HCT116 with different genetic backgrounds and degrees of intrinsic radiosensitivity (Fig. S4A). Measurement of cell viability 5-6 days post irradiation and in the presence of different concentrations of trametinib showed enhanced antineoplastic effects with combination therapy in all three cell lines. The level of radio-enhancement differed between cell lines, but could be commonly observed at low nanomolar concentrations of trametinib. Again, comparing the observed combination response with the expected combination response according to a Bliss independence model, revealed a clear excess over the Bliss model in all cell lines (Fig. 3D). In long-term colony forming assays (10-12 days), the enhancing effect of combined MEKi and radiation was also clearly visible (Fig. 3E). We also observed sensitizing effects for two pharmacological inhibitors of the RAS-MAPK pathway that were not included in the drug screening libraries, KRAS:SOS1 and ERK inhibitors (Fig. S4B-D). Compared to MEK inhibitors, the radio-enhancing effect of the two compounds was weaker and more cell-line dependent. When compared to the MDM2 inhibitor and previously reported radiosensitizer nutlin-3a,^18^ MEKi could achieve similar sensitizing effects, but at much lower drug concentrations (Fig. S4E-F) in TP53 wild-type HCT116 cells. Of note, nutlin-3a also showed minor radiosensitizing effects in our organoid assays, as compared to MEK inhibitors (Fig. S4G). These results indicate that targeting aberrant RAS-MAPK signaling by MEK-inhibitors or other inhibitors of the pathway significantly increases cellular response to radiation in CRC.

**Fig. 3:**
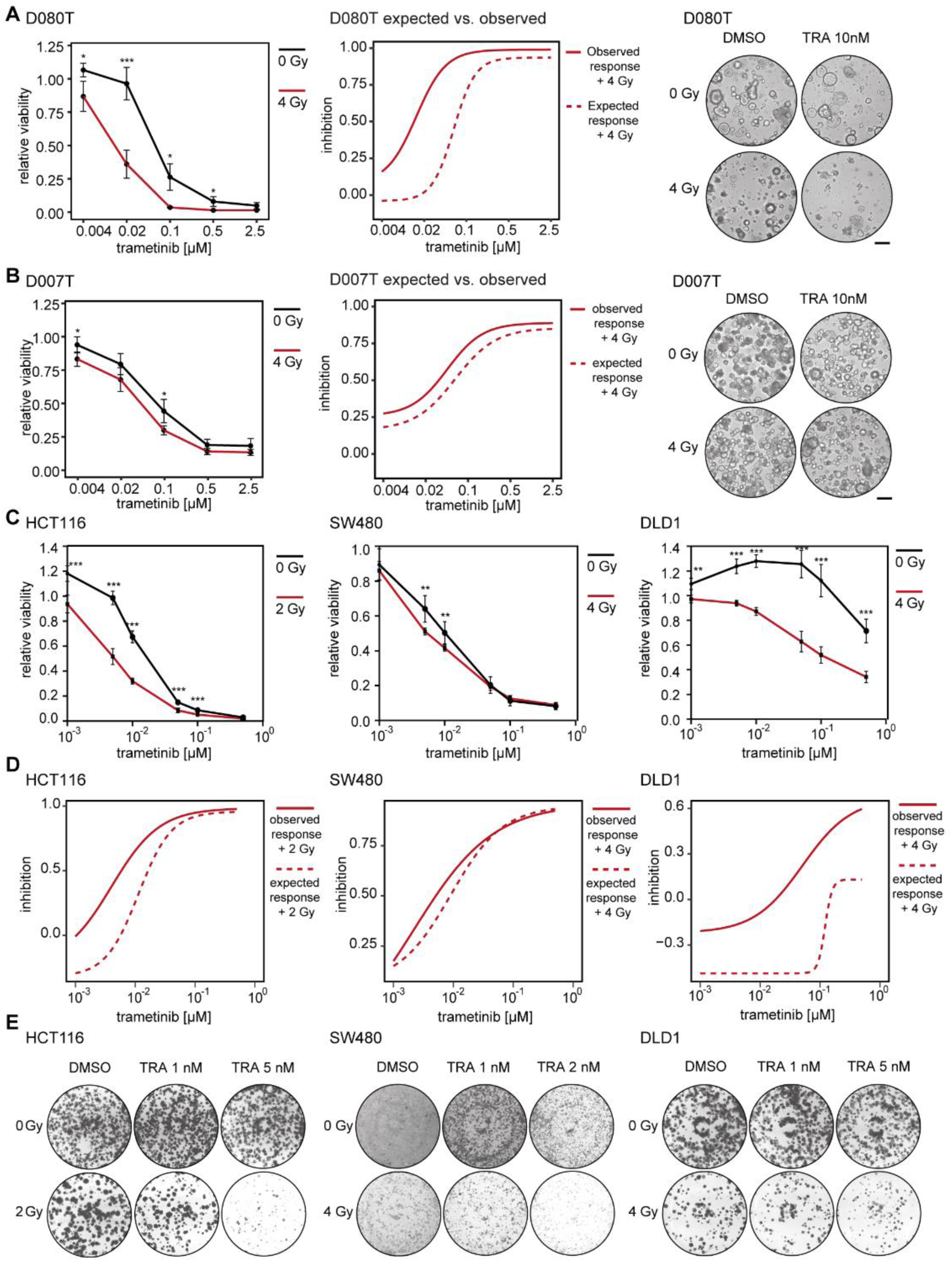
MEK inhibition is synergistic with radiation in colorectal cancer cell lines and organoids. **A,** Left: viability assay of organoid line D080T treated with increasing concentrations of MEK inhibitor trametinib with- and without radiation, data of the irradiated and non-irradiated plates were normalized to DMSO controls on the same plates in this analysis, respectively, to visualize the additional effect of trametinib in the irradiated condition. Middle: Dose-inhibition relationship of the same data. A Bliss’ independence model of trametinib and radiation was used for each concentration to calculate the expected inhibition. For this analysis, all treatments were normalized to non-irradiated DMSO controls. Right: example images of organoids, scale bar: 50 µm. **B,** Left: viability assay of organoid line D007T treated with increasing concentrations of MEK inhibitor trametinib with- and without radiation, data of the irradiated and non-irradiated plates were normalized to DMSO controls on the same plates in this analysis, respectively, to visualize the additional effect of trametinib in irradiated condition. Middle: Dose-inhibition relationship of the same data. A Bliss’ independence model of trametinib and radiation was used for each concentration to calculate the expected inhibition. For this analysis, all treatments were normalized to non-irradiated DMSO controls. Right: example images of organoids, scale bar: 50 µm. **C,** Viability assays of CRC cell lines treated with increasing concentrations of MEK inhibitor trametinib with and without radiation. Cell viability was determined after 60 hours of treatment by CellTiter-Glo. Data of the irradiated and non-irradiated plates were normalized to DMSO controls on the same plates in this analysis, respectively, to visualize the additional effect of trametinib in irradiated condition. **D,** Dose-inhibition relationship of cell lines treated with increasing concentrations of trametinib and radiation. A Bliss’ independence model of trametinib and radiation was used for each concentration to calculate the expected inhibition. For this analysis, all treatments were normalized to non-irradiated DMSO controls. **E,** Colony forming assay of CRC cell lines treated with trametinib (TRA) with and without radiation. Scans of complete wells of standard six-well plates (9.6 cm² per well) are shown. A-B, E, representative images of at least three independent biological replicates are shown. A-D, Data from three (cell lines) and four (organoids) biological replicates are presented as mean ± SD. *p < 0.05, **p < 0.01, ***p < 0.001, two-tailed t-test.

### Radiation induces RAS-MAPK signaling in CRC cell lines and organoids

To determine mechanisms underlying the radiosensitizing effects of MEKi, we first assessed activity of RAS-MAPK signaling after irradiation by measuring pERK levels, as the pathway has previously been associated with radiation response in different tumor models.^19^ We observed that radiotherapy induced a transient increase in ERK phosphorylation in CRC cell lines (Fig. 4A). The onset of RAS-MAPK activation differed between cell lines, ranging from day 2 to 4 post irradiation (Fig. 4A). Six days after irradiation, pERK levels were equal between irradiated and non-irradiated cells. Activation of RAS-MAPK signaling was also demonstrated at the level of target genes, as irradiation increased the expression of EGR1, SPRY2 or DUSP4 in CRC cell lines (Fig. 4B) and organoids (Fig. S5A). Expression profiling of three patient-derived CRC organoid lines after irradiation with 4 Gy showed a number of differentially expressed genes, some of them related to the RAS-MAPK signaling pathway (Fig. 4C, Fig. S5B). Using pathway enrichment analysis with Molecular Signatures Database HALLMARK gene sets,^20^ we found that “KRAS SIGNALING UP” was among the significantly upregulated gene sets in irradiated organoids, in addition to expected signatures such as DNA_Repair, P53_Signaling and several inflammatory pathways (Fig. S5C). We found that concomitant treatment with trametinib potently repressed the radiation-induced increase in pERK levels and expression of target genes of RAS-MAPK signaling in CRC cell lines (Fig. 4E-F). This finding was corroborated in CRC organoids, as MEK inhibition markedly suppressed ERK phosphorylation induced by radiation (Fig. 4G). In summary, our results suggest that activation of RAS-MAPK signaling presents a cellular adaptation of CRC to radiation. Targeting the pathway with MEK inhibitors could abolish this adaptive activation, thus providing a mechanism by which the drug sensitizes CRC cells to radiation.

**Fig. 4:**
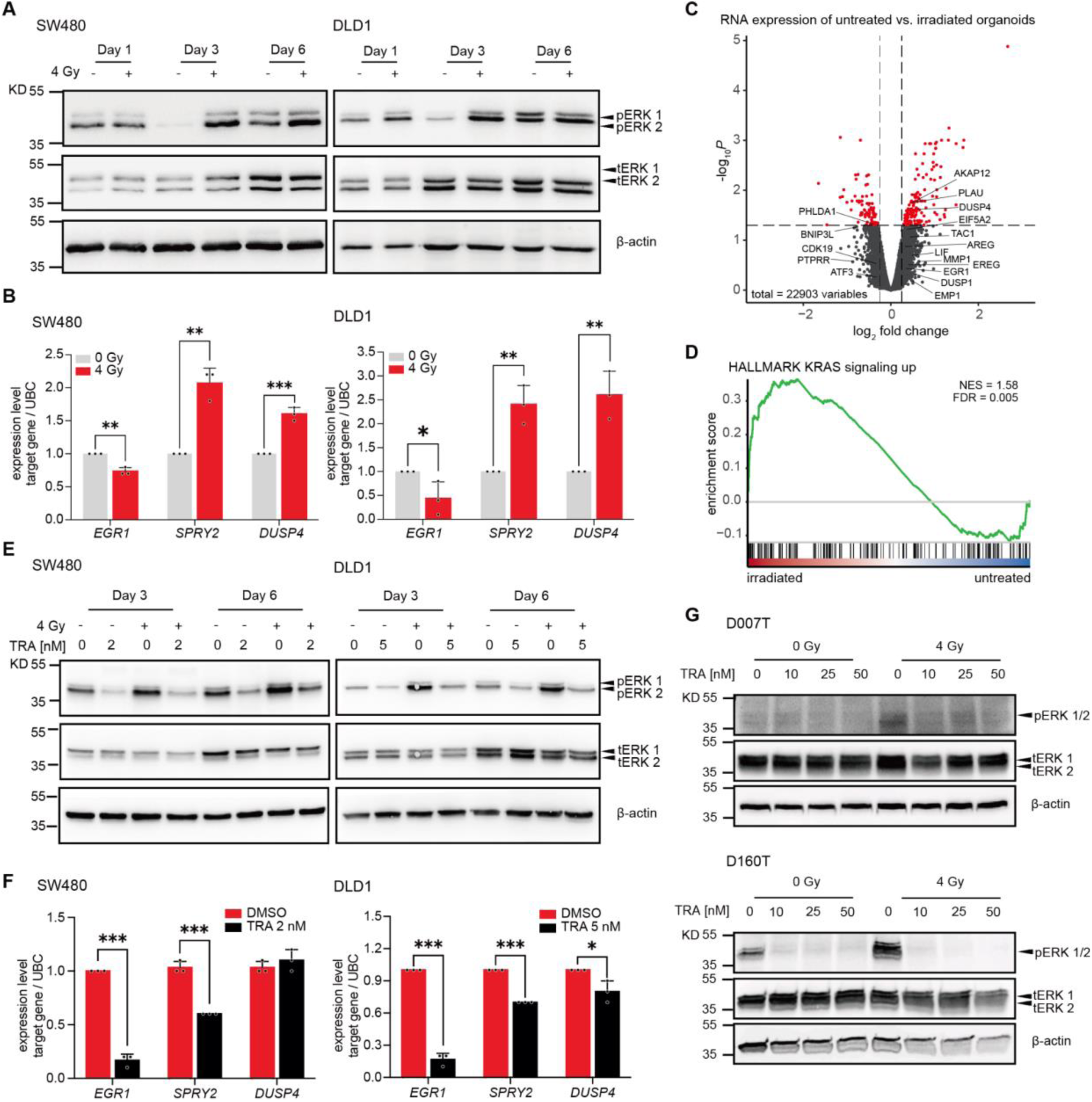
Radiation induces activation of RAS-MAPK signaling. **A,** Phosphorylation of ERK1/2 in CRC lines at different time points after irradiation. **B,** Expression of RAS-MAPK pathway target genes is induced in DLD1 and SW480 cell lines after irradiation. **C**, RNA expression profiling of rectal cancer organoid lines D007T, D080T and D160T. Volcano plot of differentially expressed genes in irradiated vs. non-irradiated organoids. Target genes of the EGFR signaling pathway according to PROGENY are highlighted. **D**, Barcode enrichment plot of HALLMARK gene set KRAS_SIGNALING_UP in irradiated vs. non-irradiated organoids. **E**, Phosphorylation of ERK1/2 in CRC lines after irradiation is reduced by MEK inhibitor trametinib (TRA) treatment. **F**, Transcriptional induction of target genes of RAS-MAPK pathway after irradiation is suppressed by MEK inhibition in CRC cell lines. **G**, Phosphorylation of ERK1/2 in rectal cancer organoids after irradiation is reduced by the MEK inhibitor trametinib. **A, E, G**, Representative images of three independent biological replicates are shown. **B, F,** Data from three independent experiments are presented as mean ± SD *p < 0.05, **p < 0.01, ***p < 0.001, two-tailed Student’s t-test.

### MEK inhibition interferes with DNA damage response via repression of RAD51

The DNA damage response pathway is activated upon radiation-induced DNA double strand breaks (DSB). We assessed if targeting MEK1/2 affects this process by first measuring the formation of DSB upon radiation and the kinetics of their resolution in the presence of the inhibitor. As shown in Fig. 5A-B, radiation rapidly caused the formation of γH2AX positive foci in the nucleus of CRC cells which decreased over time. Concomitant treatment with MEKi directly after irradiation neither caused an increase in the number of foci per nuclei nor changed the speed of their resolution (Fig. 5B). This observation was confirmed by immunoblot analysis of p-γH2AX levels, which are increased upon irradiation, but not reduced by MEKi (Fig. 5C). Next, we analyzed if subsequent steps of the DNA repair pathway are affected by MEKi. DSB can be repaired by two distinct pathways of the DNA repair machinery^21^ and we first measured transcript levels of main components of both pathways. We found that radiation upregulated the transcript levels of many DNA repair genes such as *DDB2* and *XRCC2* in CRC cell lines (Fig. S6A-B) and induced RNA expression signatures of DNA repair in CRC organoids (Fig. S6C), consistent with observations from previous studies.^22^ We then performed global proteomics profiling of the CRC cancer cell line HCT116 upon treatment with trametinib. A small set of proteins were strongly downregulated by MEKi, including RAD51, a central component of the homologous recombination DNA repair pathway (Fig. 5D).^23^ Interestingly, protein levels of other components crucial for the repair of DSB remained unchanged, including BRCA1, PARP2 or ATM (Fig. 5D, Fig. S6D). Loss of RAD51 upon MEKi was further confirmed in two CRC cell lines and two cancer organoids lines, showing a dose-dependent decrease of RAD51 (Fig. 5E-F). Interestingly, MEKi-induced reduction of RAD51 began approximately 12 h after addition of trametinib (Fig. S6E) and was not caused by a transcriptional repression (Fig. 5G). Co-treatment with two proteasomal inhibitors did not rescue MEKi-induced loss of RAD51, indicating that MEKi elicits proteasome-independent mechanisms to reduce RAD51 levels (Fig. S6F). We also observed that radiation itself increased RAD51 protein levels in CRC cell lines at late time points (day 6) (Fig. S6G). Hence, we hypothesized that functional depletion of RAD51 would increase radiosensitivity in our CRC models. To this end, we used RNAi to efficiently knock-down RAD51 in CRC cell lines, which resulted in a clear radio-enhancement in colony forming assays (Fig 5H). Moreover, we used the RAD51 inhibitor RI-1 to pharmacologically target the protein function in CRC cell lines and organoids. Similar to the RNAi-mediated knockdown, RI-1 sensitized both tumor models to irradiation (Fig. 5I-J). In summary, these results indicate that MEKi-induced loss of RAD51 is a central mechanism explaining its effect as a radiosensitizer.

**Fig. 5:**
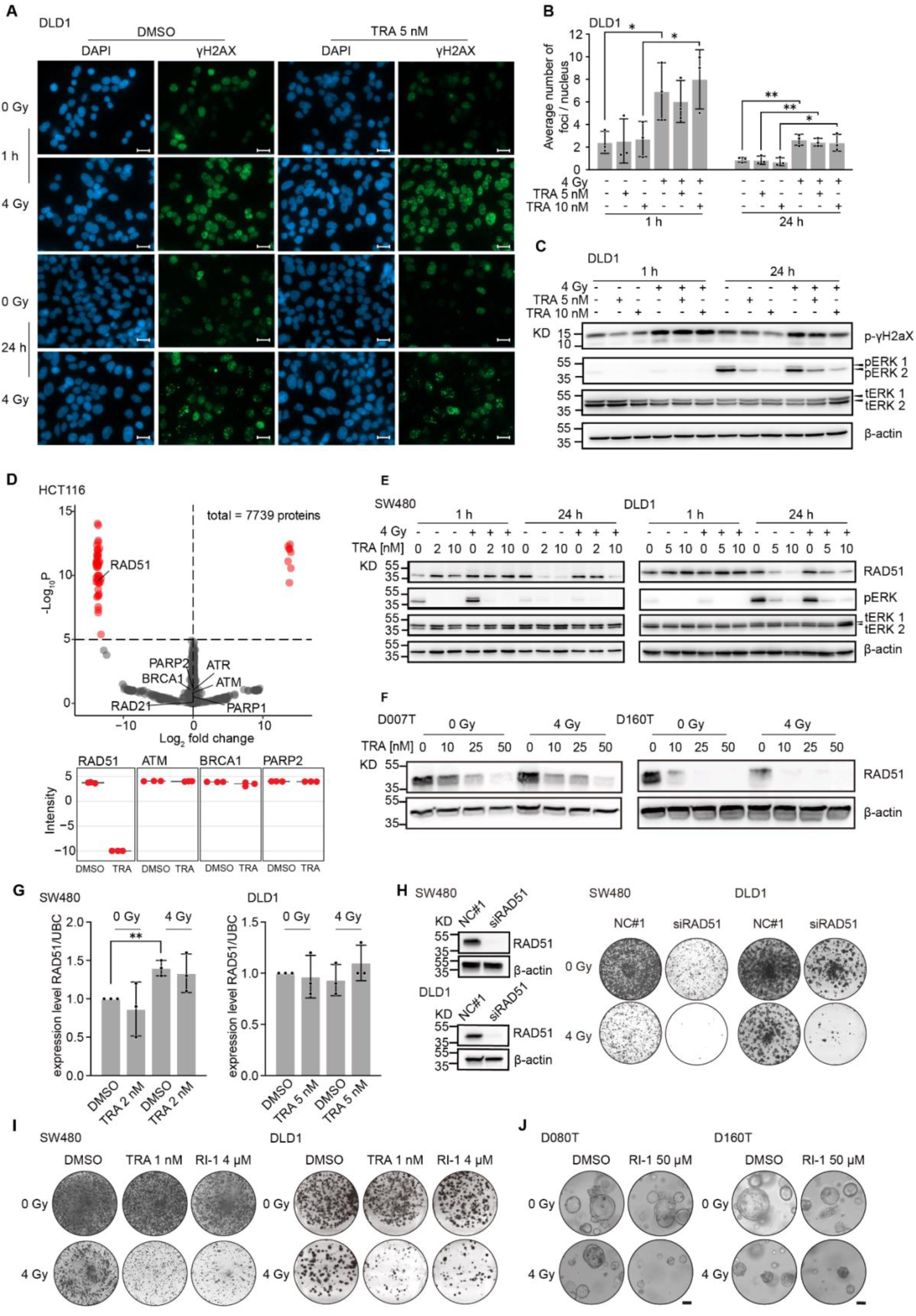
MEKi modulates DNA damage response by downregulating DNA repair gene RAD51. **A,** Radiation-induced DNA damage as determined by immunofluorescence staining of p-γH2AX. Green, p-γH2AX; blue, DAPI; 63X magnification; Scale bar: 20 µm. **B,** Measurement of p-γH2AX foci per nuclei under different treatment conditions. **C**, Immunoblot showing induction of cellular p-γH2AX levels upon radiation. **D,** Global proteome profiling by mass spectrometry of HCT116 cells after treatment with trametinib vs. DMSO for 24 h, abundance of selected DNA damage response pathway proteins is depicted below. **E**, RAD51 protein expression at different time points after irradiation and MEKi trametinib treatment in CRC cell lines. **F,** RAD51 protein expression after irradiation and MEKi trametinib treatment in patient-derived rectal cancer organoids. **G,** RNA expression levels of RAD51 in CRC cell lines after irradiation +/-trametinib treatment as determined by qPCR. **H**, Colony forming assay with CRC cell lines after siRNA mediated knockdown of RAD51 +/-radiation. Knockdown efficiency in protein expression levels is shown by western blot (left). **I,** Colony forming assay in CRC cell lines after treatment with different concentrations of the RAD51 inhibitor RI-1 with and without radiation. Scans of complete wells of standard six-well plates are shown (9.6 cm² per well) (**H,I**). **J,** Proliferation of patient-derived rectal cancer organoids after treatment with RI-1 and with / without radiation, scale bar: 50µm. **A, E, F,** representative images of three independent biological replicates are shown. **B, G,** Data from three independent experiments are presented as mean ± SD. *p < 0.05, **p < 0.01, two-tailed t-test, p-values are only shown in case of significant differences.

### MEK and PARP inhibition have synergistic effects on viability in CRC models

Besides MEK inhibitors, we also identified inhibitors of additional pathways that strongly enhanced sensitivity to radiation response in our primary screen in rectal cancer organoids. We hypothesized that a combination of MEKi with one of these inhibitors could potentiate the antineoplastic effects, particularly when combined with radiation. Specifically, combinations of MEK inhibitors with EGFR inhibitors and PI3K inhibitors were previously shown to elicit synergistic antineoplastic effects.^24^ Therefore, we performed drug combination experiments in four rectal cancer organoid lines (all of them RAS mutated, two *TP53* WT, two *TP53* mutated) using high-resolution drug concentration matrices. To this end, MEKi was combined with four drugs representing pathways with strong positive interaction with radiation (PI3K, PARP, EGFR, CHK1), under irradiated and non-irradiated conditions (Fig. 6A, Fig. S8). Focusing on drug synergy in the absence of radiation first, we determined most relevant combinations, again using the Bliss independence synergy model, and additionally tested further commonly used synergy models (highest single agent [HSA], Loewe synergy model and zero interaction potency model [ZIP]). We found that the combination of MEK inhibitor with the PARP inhibitor talazoparib and PI3K inhibitor taselisib were consistently synergistic across the tested organoid lines, while EGFR inhibitor dacomitinib and CHK1 inhibitor MK-8776 showed less consistent effects (Fig. 6B). Thus, our assay confirmed previously observed synergies of MEK inhibition and PI3K inhibition, as well as EGFR inhibition in CRC models.^24^ Since we had previously shown that MEK inhibition interferes with DNA damage response via RAD51, we further focused on PARP inhibition as a new combination partner that converged on the DNA repair pathway. We performed in-depth evaluation of combinations of MEK and PARP inhibitors at different concentrations for drug synergism. Using the Bliss synergy model, we found that the drug combination was most synergistic in lower concentrations of both PARP and MEK inhibitors, particularly in the range of 0.039 µM - 0.16 µM talazoparib and 2.4 nM - 9.8 nM trametinib in our organoid assays (Fig. 6C-F, Fig. S9). In this concentration range, the increased efficacy of the combination was clearly visible by comparing the observed response to the expected response according to the Bliss synergy model (Fig. 6G). We also confirmed these findings using two CRC cell lines, in both short- and long-term viability assays. Synergistic antineoplastic effects in short-term proliferation assays were observed in both Bliss and ZIP synergy models (maximum Bliss score: DLD1 18.68, SW480 38.41; maximum ZIP score: DLD1 15.57, SW480 44.47). Together, these results indicate that MEK and PARP inhibitors can synergistically reduce viability in different CRC models at low concentrations, even in the absence of radiation.

**Fig. 6:**
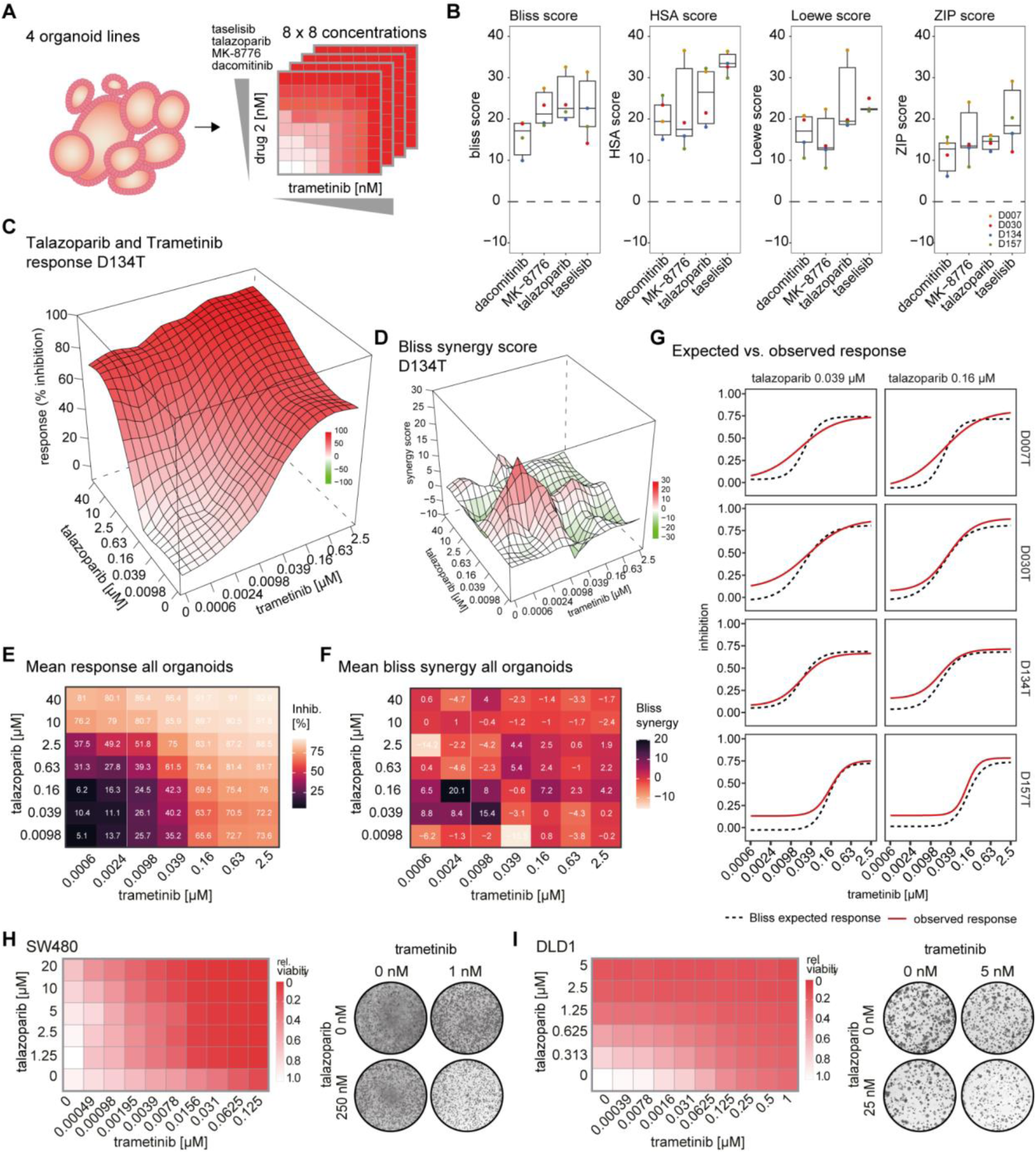
MEK and PARP inhibition have synergistic viability effects in colorectal cancer models, **A,** A combination drug screen was performed with MEK inhibitor trametinib vs. 4 other top candidates interacting with radiation, derived from the radiosensitization screening experiments shown in Figure 2 (PI3K inhibitor, PARP inhibitor, EGFR inhibitor, CHK1 inhibitor) in matrices of 7×7 concentrations (8×8 including DMSO) using 4 organoid lines. **B,** Synergy scores according to Bliss synergy, highest single agent (HSA), Loewe and zero interaction potency (ZIP) models for the 4 drugs in combination with trametinib are shown. The overall scores represent the highest score of all dose combinations tested, 2 biological replicates were analyzed for D030T and D157T, and 3 replicates were analyzed for D007T and D134T. **C,** 3-dimensional response (% inhibition) surface of the talazoparib and trametinib combination, exemplified by D134T organoids. The surface contains fitted values. **D,** Bliss synergy surface of the talazoparib and trametinib combination in D134 organoids. The surface contains fitted values. **E,** Heatmap of response (% inhibition) of trametinib-talazoparib combinations. The average of all four tested organoid lines is shown. Results of individual organoid lines are found in Fig. S8. **F,** Heatmap Bliss synergy score of trametinib-talazoparib combinations. The average of all four tested organoid lines is shown. Results of individual organoid lines are found in Supplemental Figure S8. **G,** Growth inhibition of 4 cancer organoid lines treated with increasing concentrations of trametinib in the presence of talazoparib at 0.039 and 0.16µM. Expected response according to Bliss synergy model and observed response are shown. **H-I,** viability after combinatorial inhibition of MEK and PARP in short-term and long-term viability assay in CRC cell lines. SW480 (**H**) and DLD1 (**I**) were treated for 4 days with trametinib and talazoparib in a concentration matrix, followed by cell viability measurement. Results were normalized to the DMSO control. Means of 3 biological replicates are presented (left). Long-term colony formation assays showing the combinatorial inhibition of MEK and PARP on colony growth compared to any single reagent treatment in CRC cell lines (right). Scans of complete wells of standard six-well plates are shown (9.6 cm² per well).

### Radiation synergizes with the MEK-PARP combination therapy

Having shown synergistic viability effects of MEK inhibitors with PARP inhibitors, we hypothesized that this drug combination would further synergize with radiation in CRC and allow low-dose application of both drugs in this setting, as both compounds target the DNA repair pathway. We therefore analyzed the effect of radiation on combinations of trametinib and talazoparib in cancer organoids using the high-density drug concentration matrices, as described above. In four tested organoid lines (one irradiation-resistant line, one line with stronger irradiation response, and two previously untested lines added as unbiased set), we observed a strong increase in response to combinations of the two drugs when additional radiation was performed (Fig. 7A, Fig. S9A). Applying a Bliss synergy model, we calculated the expected response of radiation added to combinations of trametinib and talazoparib and found observed responses exceeding the calculated Bliss response, particularly in the lower doses ranging from 0.6 nM to 9.8 nM trametinib combined with 0.039 µM to 0.625 µM talazoparib (Fig 7A). This proved the synergy of the two-drug combination with additional radiation. Of note, higher concentrations of the two-drug combination with radiation led to complete killing of almost all organoids, showing the high efficacy of this combination (Fig. 7A, Fig. S9A). To also prove the synergy of added PARPi to the combination of MEKi with radiation, which we had shown to be synergistic above, we calculated a second Bliss model. This model considered trametinib-radiation and added talazoparib as independent perturbations (Fig. 7B-C). The observed combination response showed higher potency than the predicted response according to the Bliss model, proving synergy between PARPi and MEKi-radiation. Again, this effect was visible mainly with the lower talazoparib concentrations between 0.039µM - 0.625µM in our assay, likely due the high efficacy of the triple combination (Fig. 7C). We also confirmed the markedly enhanced effect of the two-drug-radiation combination by long-term colony forming assays and short-term viability assays in CRC cell lines (Fig. 7D, Fig. S9B-C). Together, these findings revealed the synergistic effect of radiation with MEK inhibitor and PARP inhibitor treatment by blocking radiation-induced RAS-MAPK signaling, as well as inhibition of DNA damage response through inhibition of RAD51 by MEK inhibitors and additional targeting of this pathway with PARP inhibition (Fig. 7E).

**Fig. 7:**
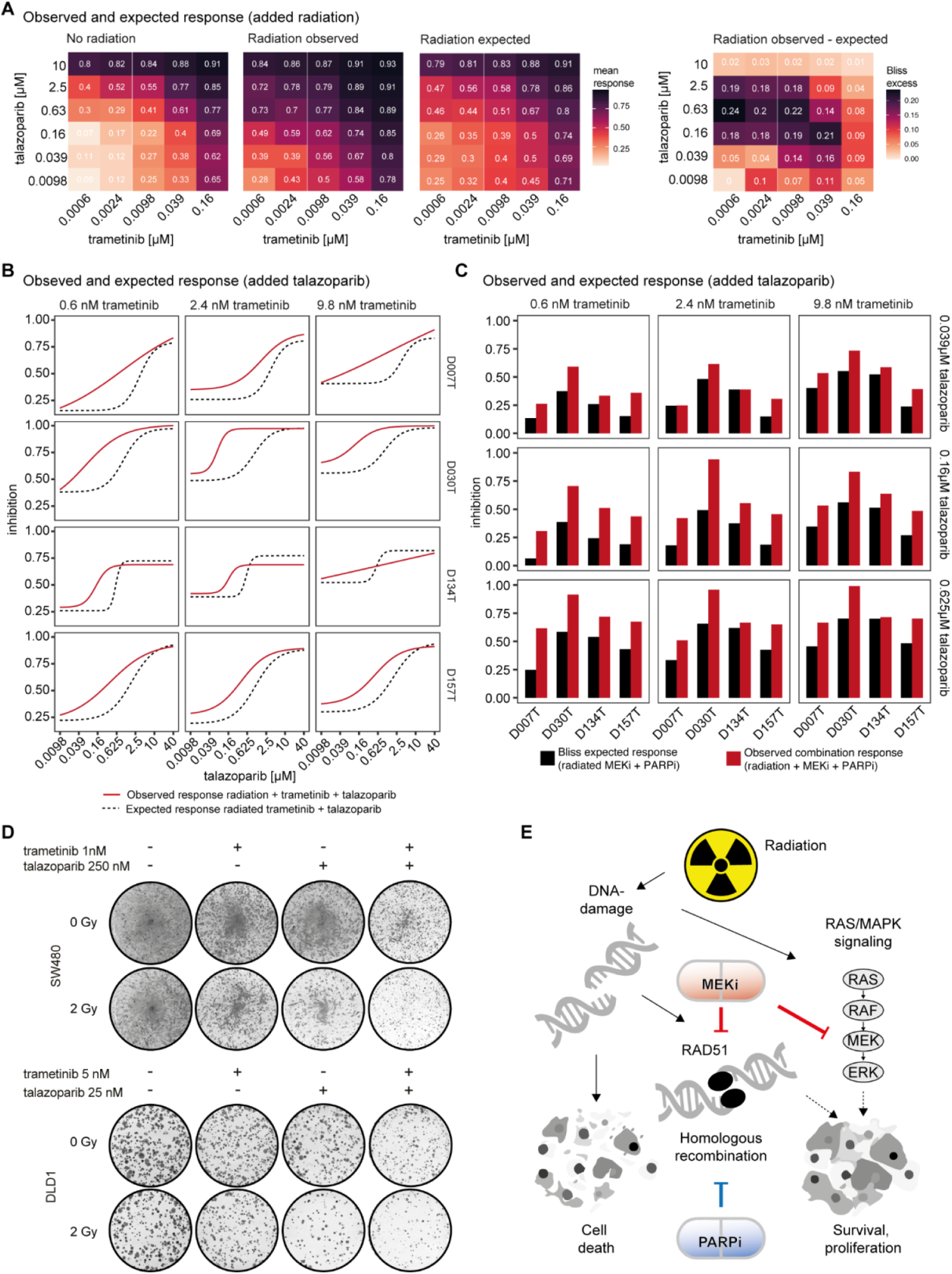
Radiation synergizes with the MAK-PARP combination therapy. **A,** Response/inhibition matrix derived from talazoparib - trametinib combinations, averaged over all four tested organoid lines: Non-irradiated, irradiated, Bliss expected response, according to a model of added radiation to fixed combinations of trametinib and talazoparib, as well as Bliss excess (observed response - expected response). Data were normalized to non-irradiated DMSO controls. The lowest 5-6 concentrations tested are shown for each drug. **B,** Dose-inhibition relationships of trametinib-radiation in combination with talazoparib treatments. Bliss’ expected response was calculated by using trametinib-radiation as one perturbation and adding talazoparib as second perturbation. D007, D030 and D157 were irradiated with 4 Gy, D134 as a more radiation-sensitive line was irradiated with 2 Gy. **C**, Bar plots of Bliss expected response vs. observed response in organoid lines treated with radiation, 0.039 - 0.625 µM talazoparib and 0.6 - 9.8 nM trametinib. Bliss expected response was calculated according to the same model as described in B. **D,** Long-term colony formation assays of radiation in combination with MEK inhibition and PARP inhibition on colony growth compared to any single reagent treatment in CRC cell lines. Scans of complete wells of standard six-well plates are shown (9.6 cm² per well). **E,** Putative mechanisms of interaction of MEK-PARP-radiation combination therapy. Radiation leads to DNA damage, which induces DNA damage repair to allow cancer cell survival. Additionally, the RAS-MAPK pathway is upregulated. MEK inhibitors block radiation-induced RAS-MAPK signaling, and downregulate RAD51, a core protein of the DNA double strand break homologous recombination repair pathway. Addition of PARP inhibitors further enhances the effect by targeting DNA damage response via a different target.

## Discussion

Chemoradiation is the main therapy for locally advanced rectal cancers, but until now, strategies that specifically target altered signaling pathways in this tumor entity are not established. Our rectal cancer organoid assays can recapitulate clinical responses to chemoradiation similar to previous reports.^10–12^ Building on this, we exploited the predictive value of our cancer organoid translational platform to discover novel drug combinations to enhance radiation response. We performed large-scale radiation-drug screens using a rectal cancer organoid platform and showed that targeting RAS-MAPK signaling by clinically approved MEK inhibitors resulted in enhanced response when combined with radiation therapy. Using different CRC models, we revealed that suppression of radiation-induced RAS-MAPK pathway activation and homology-directed DNA repair via RAD51 are major mechanisms by which MEK inhibition enhances radiotherapy. Finally, combinatorial drug-pair plus radiation experiments revealed that the effect of MEK inhibitors and radiation in rectal cancer models could be further increased by additional PARP inhibition. Thus, our study provides strong experimental rationale to combine radiotherapy with two clinically approved targeted therapies as a novel treatment strategy for rectal cancers.

A modulating effect of the RAS-MAPK pathway on radiosensitivity has been described in other cancer entities.^25^ MEK inhibitors were reported to boost the effect of radiation in pancreatic,^26^ lung^27^ and mammary cancer cell lines.^28^ Several underlying mechanisms for the radiosensitizing effect of MEK inhibitors have been described, most involving DNA damage repair. These include suppression of homologous recombination genes such as DNA-PKcs in different tumor entities,^26,29^ resulting, for instance, in a BRCA-like state in melanoma.^30^ According to our data, MEK inhibition does not affect the formation or resolution of DNA double strand breaks in CRC models, as observed in lung and pancreatic cancer cell lines.^27^ Instead, our findings suggest that RAD51, but not other important components of the DNA repair pathway, is downregulated in both CRC cell lines and organoids by MEK inhibition. RAD51 is a major component of the homologous recombination repair system and has been considered as a potential target to enhance radiosensitivity.^31^ RAD51 was also shown to be a marker of resistance to PARP inhibitors in BRCA mutated breast cancer^32^ and depletion of RAD51 via RNAi could re-sensitize cancer cells to PARP inhibition.^33^ A synergy between PARP and MEK inhibitors was observed in pancreatic and ovarian cancer, with mechanistic convergence on the HR repair pathway.^34^ Furthermore, a dual PARP-RAD51 inhibitor was developed^35^ and showed antineoplastic effects in the absence of radiation. These studies support our observation that downregulation of RAD51 is a potential mechanism for both the radiosensitizing effect of MEKi and its synergy with PARP inhibitors.

We also observed that MEKi abolished radiation-induced activation of RAS-MAPK signaling as an additional mechanism of radiosensitization. Radiation-induced DNA damage can activate RAS-MAPK signaling in untransformed cells such as fibroblasts^36^ and keratinocytes,^37^ but also in pancreatic or breast cancer cell lines.^38,39^ The underlying mechanisms described so far are manifold and include activation of ERK via GADD45β in breast cancer or stimulation of the pathway at the receptor levels via HER1/EGFR.^40–42^ ERK1/2 signaling is essential for activation of the G2/M cell cycle checkpoint in response to DNA damage by radiation^43^ and also associated with transcriptional upregulation of DNA repair genes, such as ERCC1 and XRCC1.^44^ Furthermore, radiation-induced ERK1/2 signaling can activate DNA-PKcs, which play a critical role in NHEJ-mediated DSB repair.^29^ These findings indicate that radiation-induced ERK signaling might represent a specific cellular adaptation to overcome DNA damage which can be pharmacologically targeted.

Both MEK and PARP inhibitors have been tested separately in early clinical studies as enhancers and sensitizers of radiation therapy in rectal cancers. A phase I trial has been conducted in rectal cancer patients to determine the maximum tolerated dose of trametinib added to 5-FU-based chemoradiation.^45^ A pathological complete response rate of 25% was observed at the maximum tolerated dose and the treatment was overall well tolerated. Due to the single-arm design, the radiosensitizing effect could only be estimated, but it was higher than the complete response rate of 15% of a matched historical cohort. PARP inhibitors can also increase radiosensitivity of CRC cells, particularly in the setting of XRCC deficiency.^46^ A phase I clinical trial assessed the maximum tolerated dose of the PARP inhibitor veliparib combined with neoadjuvant chemoradiation with capecitabine. The combination treatment was well tolerated with no dose-limiting grade III or IV adverse effects, and achieved a pathological complete response rate of 28%.^47^ Future (clinical) studies may also reveal if a specific subgroup of rectal cancers can particularly benefit from adding MEK-PARP inhibition to radiation therapy.

In conclusion, we used an organoid platform to discover a strong synergy effect of PARP-MEK inhibitor combination with radiotherapy in rectal cancer. We provide molecular explanations for the radiosensitizing effects of MEK inhibitors, indicating a convergence of the two inhibitors on the DNA repair pathway. Given that both PARP and MEK inhibitors show promising results in phase I neoadjuvant radiation trials with low levels of toxicity, our study advocates combining both agents with radiation in future clinical trials for rectal cancer.

## Methods

### Patients and clinical data

All patients were recruited at University Hospital Mannheim, Heidelberg University, Mannheim, Germany. We included patients diagnosed with rectal cancer in this study and obtained biopsies from their primary tumors via endoscopy. Additionally, two organoid lines from patients with primary colon cancer were used in mechanistic studies. Exclusion criteria were active HIV, HBV or HCV infections. Clinical data, tumor characteristics and molecular tumor data were pseudonymized and collected in a database. The research was approved by the Medical Ethics Committee II of the Medical Faculty Mannheim, Heidelberg University (Reference no. 2014-633N-MA and 2016-607N-MA). All patients gave written informed consent before tumor biopsy was performed.

### Assessment of clinical radiation response

Magnetic resonance image-based treatment response was assessed by magnetic resonance imaging tumor regression grade (mrTRG) according to Patel et al.^48^ and by analyzing tumor length before and after treatment. MrTRG 1 refers to the absence of any tumor signal and represents complete regression, whereas mrTRG 5 refers to only tumor signal without any fibrosis event and represents no regression. Patients with mrTRG between 1 and 3 were classified as “responders”, patients with mrTRG of 4 or 5 as non-responders. MRI images were assessed by one radiologist (MFF) with longstanding experience in MRI assessment, who was blinded to organoid response. Additionally, tumor regression grade after chemoradiation was assessed by pathological assessment according to Dworak regression grading. Grade 4 represents complete response in this system, grade 0 no response. We classified tumors with grades 3 and 4 as responders a nd tumors with grades between 0 and 2 as non-responders.

### Organoid culture

Organoid cultures were extracted from tumor biopsies as reported previously.^14^ In short, biopsies were washed and digested with Liberase TH (Roche) before embedding into Matrigel (Corning) or BME (Trevigen). Advanced DMEM/F12 (Life technologies) medium with Pen/Strep, Glutamax and HEPES (basal medium) was supplemented with 100 ng/ml Noggin (Peprotech), 1 x B27 (Life technologies), 1,25 mM n-Acetyl Cysteine (Sigma), 10 mM Nicotinamide (Sigma), 50 ng/ml human EGF (Peprotech), 10 nM Gastrin (Peprotech), 500 nM A83-01 (Biocat), 10 nM Prostaglandin E2 (Santa Cruz Biotechnology), and 100 mg/ml Primocin (Invivogen). 10 µM Y-27632 (Selleck chemicals) were added for thawing and passaging. Organoids were passaged every 7–10 days and medium was refreshed every 2–3 days.

### Cell lines and culture

HCT116, SW480, and DLD1 cells were obtained from ATCC. DLD1 and SW480 cells were cultured in RPMI 1640 medium (Gibco), and HCT116 cells were cultured in McCoy’s 5A medium (Gibco). 2D cell culture media were supplemented with 10% fetal bovine serum (FBS), 1 % L-glutamine and 1% penicillin/streptomycin. Absence of mycoplasma was confirmed by regular PCR-based testing.

### DNA sequencing of cancer organoids

Hot-spot mutations in cancer-related genes were analyzed as previously described by amplicon sequencing,^14^ or by exome sequencing using DKFZ-OTP.^49,50^ Variants were annotated with ANNOVAR^51^ and only exonic or splicing mutations classified as “frameshift deletion”, “frameshift insertion”, “nonsynonymous SNV”, “stopgain” or “stoploss”, with an allele frequency >0.1 present in COSMIC in hot-spot genes APC, RAS genes, TP53 and PI3CA were considered for further analysis.

### Organoid drug-radiation combination screens

#### Cell seeding

For cell seeding, organoids were first incubated with TrypLE (Life technologies) at 37°C until small clusters and single cells were obtained. Chemical separation was supported by mechanical shearing using a 1000 µl pipette and digestion was visually controlled by light microscopy. Organoid fragments were filtered through a 40 µm strainer (pluriSelect) to prevent large organoid clusters and afterwards quantified as previously described.^14^ For seeding, the required number of organoids was resuspended in the seeding medium and 50 µl of organoid suspension were seeded into each well using a multidrop dispenser (Berthold), before centrifugation for 10 minutes at 1000g at room temperature. Culture medium was supplemented with Y-27632 and growth factor-reduced BME type 2 was added to a concentration of 0.75 mg/ml. Additional organoids were seeded in a proliferation plate running in parallel to determine the proliferation during the incubation period of radiation treatment.

#### Drugs and compound libraries

For the drug-radiation combination screen two libraries were used: A kinase inhibitor library with 224 compounds (Table S2) and a clinical library of 140 drugs of which the majority was clinically approved, supplemented with selected inhibitors of interest for enhancing radiation (Table S3). The clinical library contains a comprehensive selection of FDA-approved cancer-targeting small molecule drugs, which can be modeled in our platform. Antibodies, antibody-drug-conjugates as well as drugs mainly targeting the immune system or tumor microenvironment were excluded. The kinase library was used at a maximum concentration of 10 µM and three ten-fold dilution steps to finally screen 4 different drug concentrations. Drugs within the clinical library and their maximum concentrations were selected individually based on literature review of 2D and 3D cell culture assays, as well as own previous experiments. Each compound was screened in five different concentrations after each five-fold dilutions. Thus, a total of 1596 drug perturbations (considering all drugs and concentrations) were tested in our assays. 5 µM bortezomib was used as positive control, DMSO as negative control. Both libraries were arranged in a random layout using a Biomek NX^P^ robotic system (Beckman Coulter). All drugs were purchased from Selleck chemicals.

#### Drug-drug-radiation combinations

For testing drug-drug-radiation combinations, trametinib was used in combination with talazoparib, MK-8776, taselisib and dacomitinib. Each was used in seven concentrations; each four-fold diluted. Including DMSO, each trametinib concentration was combined with all other drugs in 8 concentrations in an 8 x 8 combination matrix. DMSO was used as negative, bortezomib in 5 µM as positive control.

#### Drug treatment

Drug treatment was performed on day 3 after seeding. Old medium was aspirated, discarded and 45 µl fresh medium was added, drugs were pre-diluted in medium and 5 µl of diluted drugs were added. All pipetting steps were performed by a Biomek FX^P^ robotic device (Beckman Coulter). Plates were covered with plastic lids for radiation treatment.

#### Radiation treatment of organoids

Radiation was performed about two hours after drug treatment. Organoids were irradiated using a MultiRAD 225 cabinet X-ray (Precision X-Ray) with a voltage of 200 kV, a current of 17.8 mA and a 0.5 mm copper filter. X-Ray dose rate was 2.151 Gy/min. The dose rate was regularly controlled and re-calibrated by the department for radiation protection and radiological dosimetry (DKFZ). After radiation, plates were sealed using PlateLoc (Agilent) and incubated for 6 days at 37°C and 5% CO_2_.

#### Viability readout

Viability was measured on day 9 after seeding. Medium was aspirated and discarded before 30 µl undiluted CellTiter-Glo (Promega) solution was added to each well. After 30 minutes of incubation at room temperature, luminescence was measured by a Mithras reader (Berthold technologies).

### Radiation treatment of cell lines

In *vitro* irradiation of cell lines was conducted using 6 MV X-rays emitted by a clinical linear accelerator (Versa HD, Elekta Synergy) at a dose rate of 6.67 Gy/min with a 40 × 40 cm^2^ irradiation field. Cells were irradiated in the cell-culture plates at source-surface distance of 100 cm while using 15 mm water-equivalent material for dose build-up and 8 cm for backscatter, as described by Veldwijk et al.^52^ The dosimetry was regularly performed by medical physicists from the Department of Radiotherapy, University Medical Center Mannheim.

### Cell line viability assay

Cells were seeded at a concentration of 2000 cells per well in 96-well plates. Twenty-four hours post seeding, cells were irradiated or sham irradiated (same experimental procedure without applying radiation). Following radiation, cells were treated with either DMSO or drug. Cellular viability was analyzed 5 to 6 days after drug treatment, depending on individual growth rates of CRC cell lines. Cellular viability was determined using CellTiter-Glo assay (Promega) according to the manufacturer’s protocol. Readout was performed using a multi-function-reader Infinity M200 (Tecan).

### Cell line colony formation assay

Cells were seeded at a concentration of 1000 to 4000 cells per well in six-well plates. Twenty-four hours post seeding, cells were irradiated. Following radiation, cells were treated with either DMSO or different concentrations of small molecule inhibitors and incubated in standard conditions of temperature and humidity for 11 d. After this time, plates were washed with PBS, fixed with a methanol and acetic acid solution and stained with 0.05% crystal violet solution (Sigma Aldrich). Plates were scanned and the complete scans of each well from standard 6-well plates (Greiner Bio-one, 9.6 cm² per well) are shown in the figures without cropping, if not otherwise specified.

### Organoid proliferation assay

Luminescence of the organoids was measured on day 3 after seeding using CellTiter-Glo assay as described above. Luminescence was compared to untreated controls of the radiation response assay to determine the doubling time of each organoid line between the day of treatment and the day of readout.

### γH2AX foci assay

Cells were seeded on coverslips in six-well plates and treated with radiation and/or trametinib. One hour and twenty-four hours post radiation, cells were fixed with 3.7% paraformaldehyde in PBS for 20 min, and then blocked with 0.5% Triton X-100 with 1% BSA in PBS for 1 h. The fixed cells were incubated with anti-γH2AX antibodies (Abcam, ab26350, dilution 1:200) for 1 h at room temperature, and then were incubated overnight at 4 °C with Alexa Fluor 488-labeled secondary antibody (Fisher Scientific, A32723, 1:200), and mounted with DAPI (Vector Laboratories). Images were acquired using a Leica Axio Observer Z1/Apotome microscope. The cell numbers in regions of interest were counted manually in DAPI-stained images with the “multi-point” tool in ImageJ. Foci counting were determined in the matched γH2AX-stained images using the “find maxima” tool in ImageJ, with a fixed prominence set and manual correction for all images under the same condition. A total of 100 cells per condition were analyzed to determine the average foci number per cell.

### Quantitative PCR

Total RNA was isolated from cells using the peqGOLD Total RNA Isolation Kit (PeqLab). cDNA was synthesized using the Verso cDNA synthesis kit (Thermo Fisher Scientific) with 1 μg of purified total RNA as input. Quantitative PCR was performed in a MicroAmp® 96-well reaction plate (Life Technologies) on a StepOne Plus Real Time PCR instrument (Thermo Fisher Scientific). UBC was used as reference gene for relative quantification. Primer sequences used for quantitative PCR are listed in Table S4.

### Immunoblot

Protein extraction was performed using RIPA lysis buffer (Thermo Fisher Scientific) supplemented with protease inhibitor tablets (Roche) and phosphatase inhibitor cocktails 1-2 (Sigma Aldrich). Protein concentration was measured by BCA protein assay (Thermo Fisher Scientific). Fifteen to thirty micrograms of lysates were separated on 4-15% precast Mini-PROTEAN® TGX™ gels (Bio Rad) and transferred to nitrocellulose membrane (Amersham). Membranes were detected by chemiluminescence staining protocol with SuperSignal™ chemiluminescent substrate (Thermo Fisher Scientific). Images were acquired using the FUSION-SL-Advance imaging system (PeqLab). All antibodies used are listed in Table S5.

### RNA interference

SW480 and DLD1 cells were seeded on six-well plates (Greiner) at a density of 1 × 105 cells per well. Twenty-four hours after seeding, cells were transfected with siRNAs (siGenome RAD51, Dharmacon and siGenome non-targeting control scrambled siRNA, Dharmacon) and Lipofectamine RNAiMAX (Thermo Fisher Scientific) with a final concentration of 5 nM siRNA per well. For subsequent expression analysis of target genes and proteins, cells were harvested 48 h post transfection. For further treatment of transfected cells, the medium containing siRNAs was removed 48 h after transfection and cells were re-seeded into six-well plates or 96-well plates. Twenty-four hours after cell re-seeding, drug and/or radiation treatment was performed.

### Mass Spectrometry (MS)

HCT116 cells were seeded on 6-well plates at a density of 20.000 cells/cm². Twenty-four hours after seeding, cells were treated with 100 nM trametinib or DMSO as control for 24 hours. Cells were then washed twice with ice-cold PBS and harvested on wet-ice using a cell scraper with 200 µl of ice-cold PBS containing protease inhibitors. The cell suspension was then pelleted by centrifugation at 1000 rpm at 4 °C. The supernatant was discarded and the cells were stored at −80 °C until further processing. For MS analysis, cell pellets were lysed in 100 µl of lysis buffer containing 6 M guanidine hydrochloride (GuHCl), 5 mM tris-(2-carboxyethyl)phosphine and 10 mM chloroacetamide, boiled for 10 min at 99 °C, briefly cooled down on ice and sonicated using a BioRuptor (Diagenode, Seraing) set to high intensity with 10 cycles (30 sec ON / 30 sec OFF) at 4°C. After sonication, the lysates were centrifuged at 15.000 g for 10 min at 4 °C and the supernatants transferred into new tubes. The volume containing the equivalent of 20 µg of total proteins from each sample was transferred into new tubes and diluted to a final concentration of maximum 2 M GuHCl with 25 mM Tris-HCl buffer pH 8.5. The proteins were then digested by trypsin (sequencing-grade, Promega) at 1:50 ratio at 37 °C overnight. After overnight incubation, the samples were acidified by adding formic acid at 1% final concentration to stop the digestion. Prior to MS analysis, a peptide clean-up procedure was performed for each sample using SP3 method, as described^53^.

The quantitative measurements were carried out using an EASY-nLC 1200 system (Thermo Fisher Scientific) coupled to a Q Exactive HF Orbitrap mass spectrometer (Thermo Fisher Scientific). The peptides were separated by reverse-phase liquid chromatography with 0.1% formic acid (solvent A) and 80% acetonitrile supplemented with 0.1% formic acid (solvent B) as mobile phases, using a stepped gradient from 4% to 80 % solvent B in 120 min on a nanoEase M/Z peptide BEH C18 column (Waters, 250mm x 75 µm 1/PK, 130 Å, 1.7 µm) heated to 55 °C using a HotSleeve+ column oven (Analytical Sales & Services). The peptides were eluted with a constant flow rate of 300 nl/min.

The Q Exactive HF Orbitrap mass spectrometer was operated in data-independent mode (DIA) with a scan range of 350-1650 m/z, orbitrap resolution 240000 FWHM, 3e6 AGC target and maximum injection time (max. IT) 20 ms for MS1 scan. For MS2 scan the parameters were set as follows: orbitrap resolution 30000 FWHM, AGC target 1e6, max. IT 40 ms and the precursors were analyzed in a sequence of 26 windows of variable width with an overlapping region of 0.5 Da from both sides. The normalized collision energy for the fragmentation of precursor ions was set to 27 and a fixed first mass of 250 m/z was set for the acquisition of the MS/MS spectra.

#### Data analysis

The files containing spectral data were analyzed using Spectronaut™ (version 18) software using a directDIA™ workflow against a nonredundant UniProt Human Proteome FASTA database from 30.01.2020 with the identification settings as follows: precursor Q-value cutoff 0.01, precursor posterior error probability (PEP) cutoff 0.2, protein Q-value cutoff (experiment-wise) 0.01, protein Q-value cutoff (run-wise) 0.05, protein PEP cutoff 0.75. For the quantification, the data were normalized based on a retention time-dependent local regression model as described,^54^ with precursor filtering based on identified Q-value, and maxLFQ quantification method based on inter-run peptide ratios. The proteins were grouped by protein group ID and peptides were grouped by a stripped peptide sequence of the identified precursors.

### RNA microarrays

Organoid RNA was isolated with the Qiagen RNeasy kit following the manufacturer’s instructions. Organoids were pelleted by centrifugation and frozen in RLT buffer containing 1% β-mercaptoethanol before RNA isolation. Samples were hybridized on Affymetrix Human Genome U133 plus 2.0 arrays. Data were analyzed as previously reported.^14^ In short, raw microarray data were normalized using the robust multi-array average (RMA) method^55^ followed by quantile normalization as implemented in the “affy” R/Bioconductor package.^56^ Differential gene expression analyses were performed using a moderated t-test as implemented in the R/Bioconductor package “limma”.^57^ Gene set enrichment analyses were performed as implemented in the “fgsea” R/Bioconductor package for ranked gene lists.^58^

### Data analysis

T-test was used to compare between two experimental conditions if not otherwise stated. Statistical analysis of cell line experiments was performed in GraphPad Prism 8.0. Analysis of large-scale drug and radiation assays with organoids was performed using R. The area under curve (AUC) analysis in the GraphPad Prism was used to determine the AUC of radiation dose-response curves in cell lines. Statistical significance was indicated with asterisks: *p < 0.05, **p < 0.01, ***p < 0.001. Error bars represent standard deviation (SD) of multiple biological replicates as denoted in the figure legends.

#### Quality controls in drug-radiation screens

Pearson correlation between replicates, as well as descriptive statistics of positive and negative controls were calculated. The coefficient of variation (CV) was calculated using the standard deviation of negative controls (sd(-)) and the mean of negative controls (mean(-)) as a measurement of the negative controls‘ distribution.

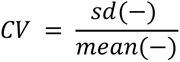

If possible, Z‘-Factor was calculated as an additional parameter for the distribution of positive and negative controls in drug-radiation screens. For its calculation the standard deviation of positive (sd(+)) and negative controls (sd(-) as well as the mean of positive (mean(+)) and negative controls (mean(-)) were used.

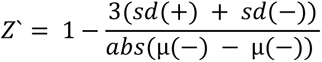

Replicates with a Z‘-Factor < 0.25 or a CV > 0.25 were excluded from analysis.^17,59^

#### Dose-Response Curves and Area under the curve

Raw luminescence data were normalized to the mean of the radiation-specific DMSO-controls platewise. Relative viability values were plotted against the radiation doses to obtain dose-response curves. To obtain comparable AUC values in drug-radiation assays, the logarithmic breaks were projected on a linear axis with the highest concentration being projected to the value 1 and the lowest concentration to the value 0. The other concentration values were equally distributed between 0 and 1 to obtain uniform breaks. Area under the curve was calculated using trapezoid integration implied in the pracma package in R (https://cran.r-project.org/web/packages/pracma/index.html).

#### Doubling time calculation

Doubling Time (Td) was calculated during the incubation period after radiation treatment. The starting point was the day of treatment (day 3 = t_1_) and the end point was the day of readout (day 9 = t_2_). Additionally, the luminescence values of treatment day (lum_1_) and readout day (lum_2_) were used for calculation.

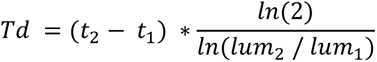

#### Growth rate inhibition metrics

For growth rate adjusted response analysis of the organoids‘ response to radiation, the method by Hafner et al.^60^ was used, to calculate the growth rate inhibition (GR) for each radiation dose (d). The luminescence at the day of treatment (lum_0_) and the luminescence at the day of readout after treatment with the respective dose (lum_d_) as well as the luminescence without treatment (lum_ctrl_) were used for calculation.

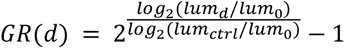

For the radiation and drug combination screens, viability was calculated by dividing the luminescence of each condition by the mean luminescence of DMSO controls. The drug concentration was plotted on a logarithmic x-axis. For AUC calculation based on drug concentration, the x axis was divided by uniform breaks between 0 and 1 as described above, while the radiation dose has a linear format.

#### Drug-drug and drug-radiation combination data analysis

Analysis of drug combinations in different organoid lines were done by calculating Bliss synergy to determine excess over the Bliss model as marker for Synergy, similar to recently published work.^17^ For Bliss excess, “the single-agent activities of drug A and drug B must be expressed as a probability between 0 and 1

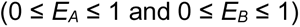

The observed effect of the combination is also expressed as a probability:

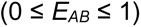

This means that the expected Bliss additive effect can be expressed as:

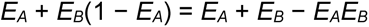

A positive “excess” over the expected Bliss additive effect defines a synergistic response”.^17^ Expected responses according to the Bliss model were compared to the observed response at each dose to identify synergistic dose regions. For synergy calculation of drug-radiation combinations, the Bliss model was used in a similar way to obtain expected response for each drug dose with radiation combination, each normalized to non-irradiated DMSO controls. To analyze synergy of an additional perturbation to an established combination (i.e. radiation added to MEKi + PARPi and PARPi added to radiation + MEKi), we treated the previously established combinations as one factor and the added perturbation as the second factor in the Bliss formula:

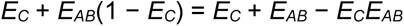

The drug-drug combinations in high density drug concentration matrices were further analyzed with the SynergyFinder Plus package to test further synergy models and visualize inhibition and synergy scores in surface plots.^61^ Growth inhibition [%] as well as its standard error (SE) were estimated by bootstrapping of the included replicates. For D030T and D157T, 2 biological replicates were analyzed and 3 replicates for D007T and D134T. Four different synergy scores were calculated for each drug combination: The Bliss model, as described above, the highest single agent (HSA), Loewe and Zero interaction potency (ZIP) score as implemented in the SynergyFinder Plus package. Metrics are reported as the highest value found across the entire dose matrices to enable identification of dose-specific maxima of synergy that may be “canceled out” when considering the average values of the full dose matrix, as previously reported.^17^

### Statistics and reproducibility

The sample size (n), replication and statistical test used for each experiment are specified in the figure legends and methods for each experiment. Power calculations were not performed to determine the sample size before each experiment. Sample sizes were chosen on the basis of experience with the given experiments.^14,62,63^ Two-tailed unpaired Student’s t-test or Welch’s t-test were used to analyze statistical significance between two groups. Statistical analyses were performed using R v 4.4.0 or GraphPad Prism v 8.0. P values <0.05 were considered as statistically significant. Two rectal cancer organoids established for radiation testing were excluded from analyses and further experiments due to potential cross-contamination. One plate of D030T and of D134T in the drug-drug-radiation combination profiling experiment, as well as one replicate of D080T in the drug-radiation-screen with the clinical drug library were excluded from further analysis due to exceeding Z‘-factor or CV cut-offs. Clinical response evaluation was blinded to organoid radiation response. Data collection and outcome assessment for other experiments were not blinded.

## Acknowledgements

We thank the NGS Core Facility of the German Cancer Research Center for help with exome sequencing and the Omics-IT Facility of DKFZ for help with sequencing data analysis. We thank the DKFZ Microarray core facility for performing expression profiling experiments. We thank Rosemarie Euler-Lange, Miriam Bierbaum and Adriana Grbenicek for their help with radiation experiments with organoids and cells. We thank Dr. Junyan Lu for discussions on data analysis. This project was supported by Seed funding to the project “RASCAL” of the DKFZ Hector Cancer Institute at University Medical Center Mannheim. QX and LW were supported by a scholarship of the Chinese Scholarship Council. JER was supported by a scholarship of the German Academic Scholarship Foundation. ZL was supported by the oversea study program of the Guangzhou Elite Project. PA was supported by funding from the DKFZ International PhD Program. RI, NV, JK, EE and TZ were supported by SFB1324. MPE is supported by the DFG (GRK 2727, B1.3.). Research in the Laboratory of JB is supported by the Hector Foundation II.

## Author contributions

Conceptualization: QX, JER, TZ, JBe. Collection of data: QX, JER, TMu, ZL, JBu, PA, ML, NV, OS, AK, EE, LW, SB, NS, DS, KEB, YP, TMi, EB, CH, MV, CBD, RI, JK, IK. Data Analysis and interpretation: QX, JER, TMu, PA, EV, MB, MPE, TZ, JBe. Drafting the manuscript: QX, JER, TM, TZ, JB. Contributions to the manuscript: ZL, JB, PA, ML, NV, OS, AK, KEB, YP, TMi, EB, CH, MV, CBD, RI, JK, IK, MB, MPE. Final approval of the manuscript: All authors.

