## Supplemental Information for "An organoid platform reveals MEK-PARP co-targeting to enhance radiation response in rectal cancer"

#### Supplementary Tables

##### Table S1: Patient characteristics

| Organoid | Sex | Location | Biopsy | T | N | M | Stage (UICC) | Grading (WHO) | Neoadjuvant Treatment | Dworak | mrTRG | Δ tumor length |
|---|---|---|---|---|---|---|---|---|---|---|---|---|
| D004T | f | rectum | primary | 3 | 2 | 0 | 3 | 2 | 50,4 Gy + capecitabine | 1 | 3 | -2 |
| D007T | m | rectum | primary | 3 | 3 | 0 | 3 | 2 | 50,4 Gy + capecitabine | 2 | 3 | 6 |
| D027T | m | rectum | primary | 4 | 1 | 1 | 4 | 2 | NA | NA | NA | NA |
| D030T | f | colon | primary | 3 | 0 | 0 | 2 | 2 | NA | NA | NA | NA |
| D046T | f | rectum | primary | 2 | 1 | 0 | 3 | 2 | 50,4 Gy + capecitabine | 4 | 1 | -6 |
| D052T | m | rectum | primary | 3 | 0 | 0 | 2 | 2 | 50,4 Gy + capecitabine | 2 | 5 | 0 |
| D073T | m | rectum | primary | 3 | + | 0 | 3 | NA | 50,4 Gy, FOLFOX | 1 | 3 | NA |
| D080T | f | rectum | primary | 3 | 2 | 0 | 3 | 1 | 50,4 Gy + capecitabine | 1 | 5 | 1 |
| D082T | f | rectum | primary | 4 | 2 | 0 | 3 | 2 | 50,4 Gy + capecitabine | 1 | 4 | -1 |
| D086T | f | rectum | primary | 3 | 2 | 0 | 3 | 2 | 50,4 Gy + capecitabine | 1 | 3 | -1 |
| D104T | f | rectum | primary | 3 | 2 | 1 | 4 | 2 | 5 x 5 Gy | 2 | NA | NA |
| D114T | f | rectum | primary | 3 | 1 | 0 | 3 | 2 | 50,4 Gy + capecitabine | 4 | 2 | 4 |
| D134T | f | rectum | primary | 3 | 0 | 0 | 2 | 2 | 50,4 Gy + capecitabine | 4 | 2 | -2 |
| D147T | m | rectum | metastasis | 3 | 1 | 1 | 4 | 2 | 50,4 Gy + capecitabine | 1 | 4 | -2 |
| D157T | f | colon | primary | 3 | 1 | 1 | 4 | 2 | NA | NA | NA | NA |
| D160T | m | rectum | primary | 3 | 1 | 0 | 3 | 2 | NA | NA | NA | NA |

Abbr: T, Tumor; N, node; M, Metastasis; NA, not available; Gy, Gray; +, positive; Dworak, Dworak pathological regression grade; mrTRG, MRI tumor regression grade.

**Table S4: Primers for quantitative PCR**

| Target gene | Species | Forward primer | Reverse primer |
|---|---|---|---|
| BRCA1 | human | TTGTTGATGTGGAGGAGCAA | GATTCCAGGTAAGGGGTTCC |
| BRCA2 | human | GAAAATCAAGAAAAATCCTTAAAGGCT | GTAATCGGCTCTAAAGAA ACATGATG |
| EGR1 | human | AGCCCTACGAGCACCTGAC | GGTTTGGCTGGGGTAACTG |
| DDB2 | human | CTCCTCAATGGAGGGAACAA | GTGACCACCATTCGGCTACT |
| DUSP4 | human | GGCGGCTATGAGAGGTTTTCC | TGGTCGTGTAGTGGGGTCC |
| RAD21 | human | AATTTGGCTAGCGGCCCAT | TGTCCGTAATGCCATTTTCACC |
| RAD51 | human | GGTGAAGGAAAGGCCATGTA | GGGTCTGGTGGTCTGTGTT |
| RAD51B | human | GCACAAAGGTCTGCTGATTTC | CCCATGTTGGTGGGTAATGT |
| SPRY2 | human | CCTACTGTCGTCCCAAGACCT | GGGGCTCGTGCAGAAGAAT |
| UBC | human | CTGATCAGCAGAGGTTGATCT TT | TCTGGATGTAGTCAGACAGG |
| XRCC2 | human | TCACCTGTGCATGGTGATATT | TTCCAGGCCACCTTCTGATT |

**Table S5: List of antibodies for immunoblot**

| Antibody | Company | Catalogue # | Species | Dilution |
|---|---|---|---|---|
| p44/42 MAPK (Erk1/2) | Cell Signaling | 9102 | rabbit | 1:2000 |
| Phospho-p44/42 MAPK (Erk1/2) (Thr202/Tyr204) | Cell Signaling | 4370 | rabbit | 1:2000 |
| Rad51 (D4B10) | Cell Signaling | 8875 | rabbit | 1:1000 |
| Phospho-Histone H2A.X (Ser139) | Cell Signaling | 2577 | rabbit | 1:1000 |
| β-actin (C4) HRP | Santa Cruz Biotechnologies | Sc-47778 HRP | mouse | 1:150000 |
| anti-rabbit IgG, HRP-linked | Cell Signaling | 7074 | goat | 1:5000 |
| anti-mouse IgG, HRP-linked | Cell Signaling | 7076 | horse | 1:10000 |

### Supplementary Figures

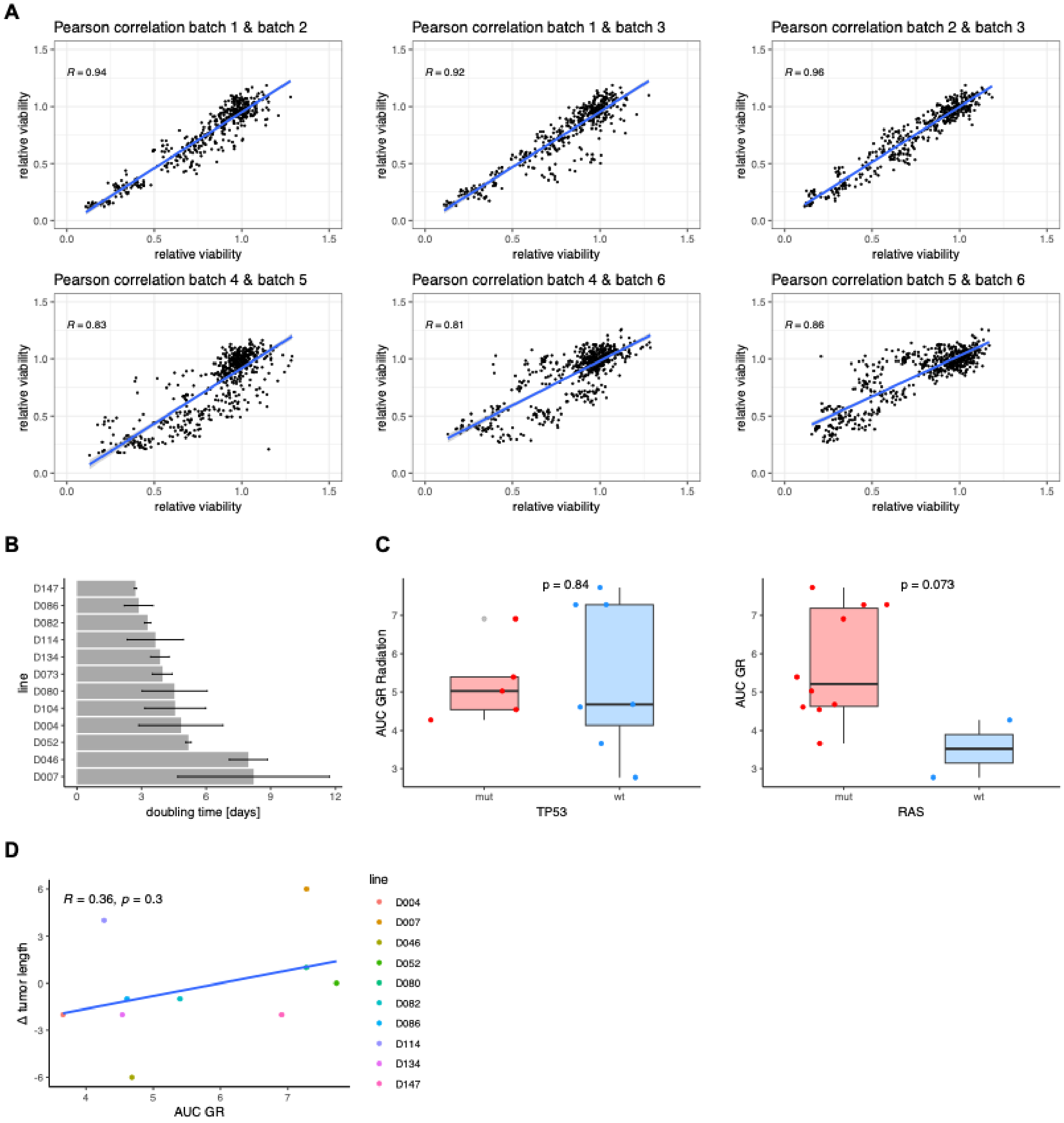

**Figure S1: An organoid platform recapitulates clinical responses of rectal cancer to radiation, related to Figure 1. A,** Quality controls for the organoid radiation assay, Pearson correlation of biological replicates of the radiation assay are shown in different batches. **B,** Doubling time of organoid lines used in the radiation response assay. Mean ± sd of three biological replicates are presented. **C,** Associations of the response to radiation and TP53 or RAS mutation status, two-tailed t-test. **D,** Pearson correlation of organoid response to radiation (AUC GR) and ∆ tumor length before and after radiation therapy, measured in MRI images.

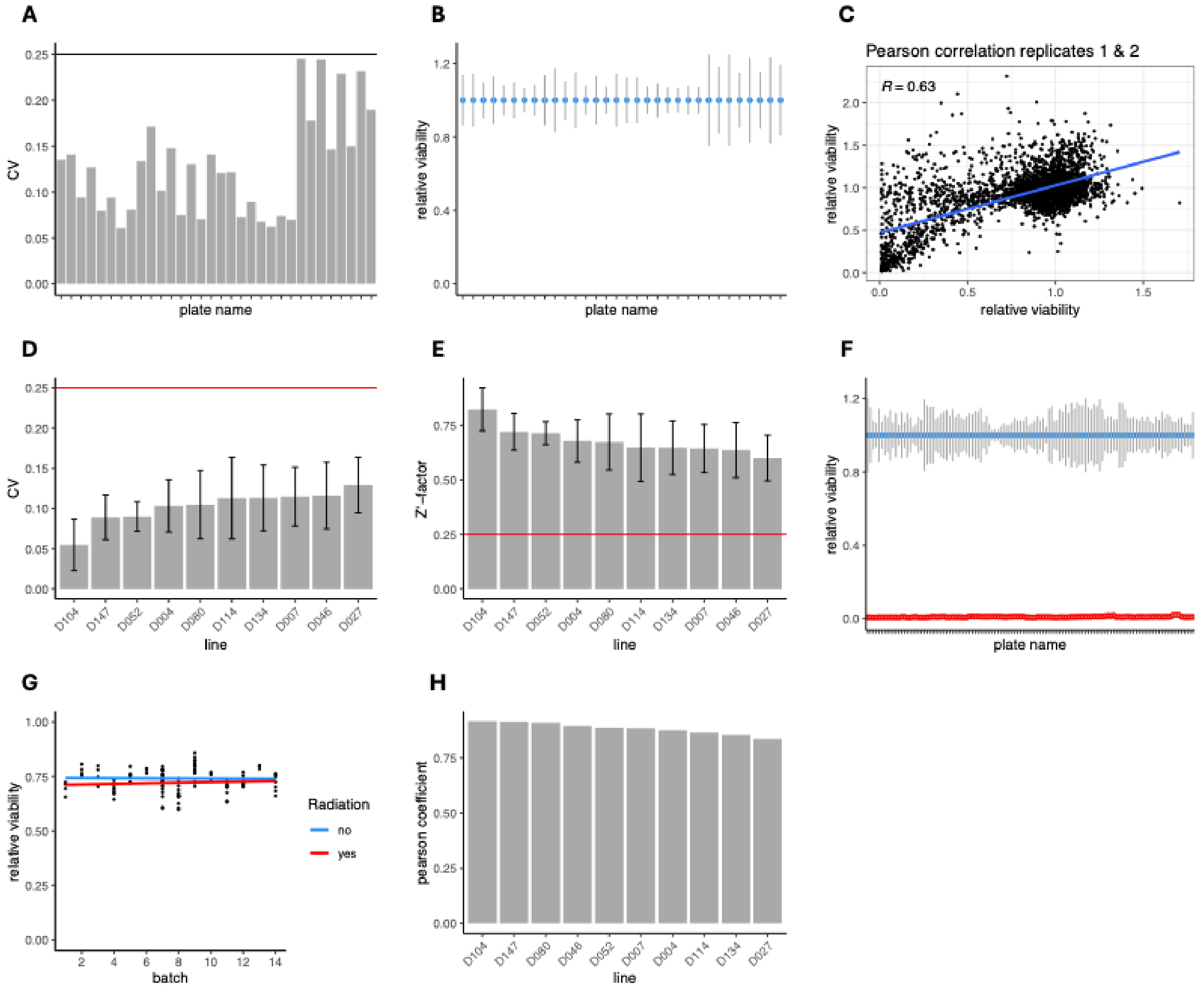

**Figure S2: Screening for synergistic effects between drugs and radiation, related to Figure 2. A-H,** Quality controls for the drug-radiation synergism screens. A kinase library of 224 drugs in 4 concentrations was tested in 2 organoid lines. 10 organoid lines were screened with a clinical drug library containing 140 compounds in 5 concentrations. DMSO was used as negative control while high-concentrated bortezomib was the positive control in the clinical library. For each line 2-4 replicates were analyzed. **A,** CV values of the DMSO controls in the kinase library combination screen are plotted for each plate. All CV values were < 2.5. **B,** Normalized luminescence values of DMSO controls for each plate are plotted as mean ± standard deviation. **C,** Pearson correlation of replicates 1 and 2 in the kinase library combination screen. **D,** Mean CV values of the DMSO controls in the clinical library combination screen are plotted as mean ± standard deviation of 2-4 biological replicates. All mean CV values were < 2.5. **E,** Z'-factor was calculated of the raw luminescence values for each plate using the formula described in the Methods section. Mean Z`-factors ± standard deviation of 2-4 biological replicates are shown. **F,** Distribution of normalized luminescence values of positive and negative controls in the clinical library combination screen. **G**, Mean viability for each plate for different batches. No tendencies were detected. **H,** Average Pearson correlation coefficient of normalized values of 2-4 biological replicates is plotted for each line.

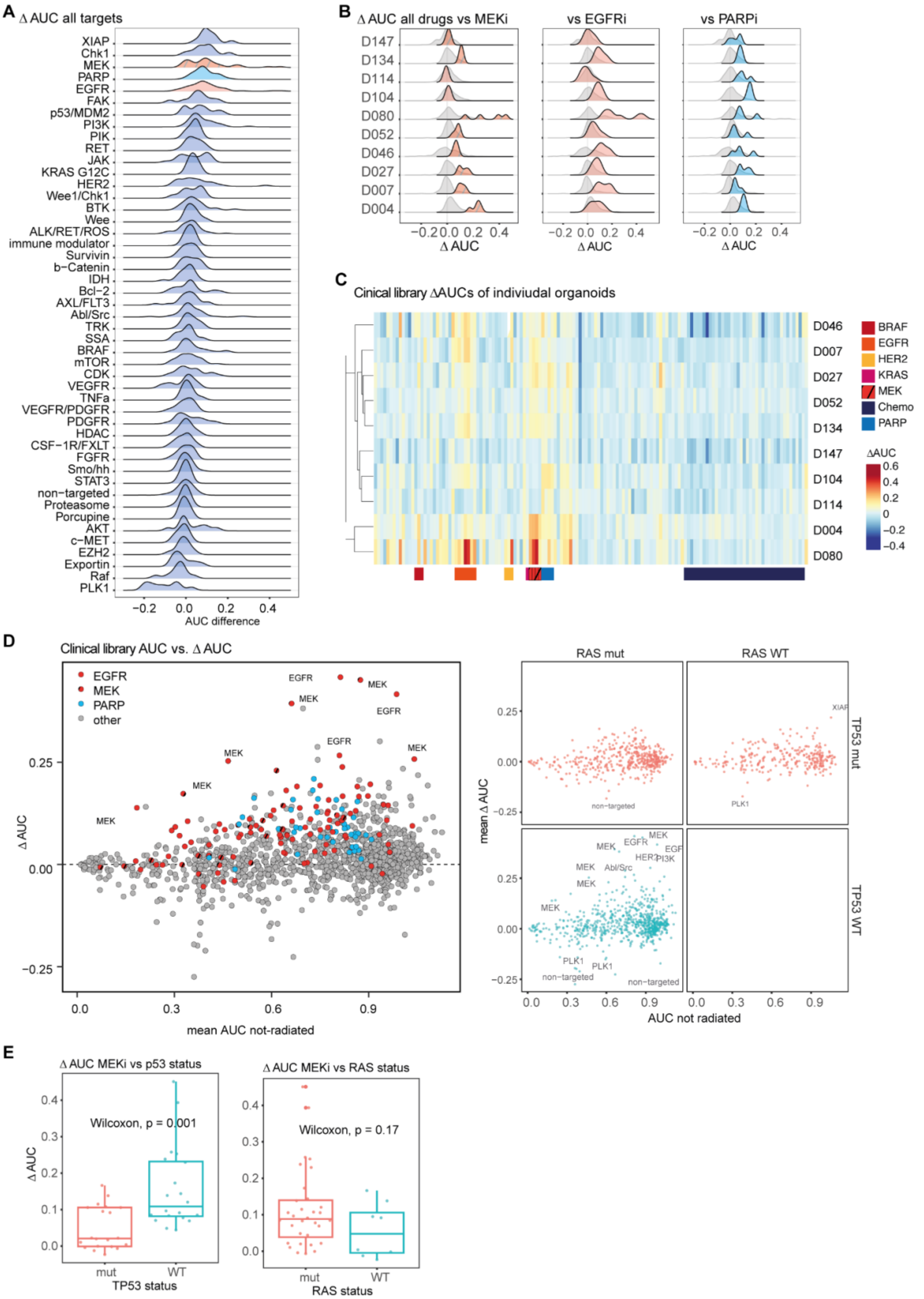

A
Δ AUC all targets
XIAP
Chk1
MEK
PARP
EGFR
FAK
p53/MDM2
PI3K
PIK
RET
JAK
KRAS G12C
HER2
Wee1/Chk1
BTK
Wee
ALK/RET/ROS
immune modulator
Survivin
b−Catenin
IDH
Bcl−2
AXL/FLT3
Abl/Src
TRK
SSA
BRAF
mTOR
CDK
VEGFR
TNFa
VEGFR/PDGFR
PDGFR
HDAC
CSF−1R/FXLT
FGFR
Smo/hh
STAT3
non−targeted
Proteasome
Porcupine
AKT
c−MET
EZH2
Exportin
Raf
PLK1
−0.2 0.0 0.2 0.4
AUC difference
B
Δ AUC all drugs vs MEKi
vs EGFRi
vs PARPi
D147
D134
D114
D104
D080
D052
D046
D027
D007
D004
−0.2 0.0 0.2 0.4
Δ AUC
C
Cinical library ΔAUCs of indiviudal organoids
D046
D007
D027
D052
D134
D147
D104
D114
D004
D080
BRAF
EGFR
HER2
KRAS
MEK
Chemo
PARP
ΔAUC
0.6
0.4
0.2
0
−0.2
−0.4
D
Clinical library AUC vs. Δ AUC
EGFR
MEK
PARP
other
Δ AUC
mean AUC not-radiated
RAS mut
RAS WT
TP53 mut
TP53 WT
mean Δ AUC
AUC not radiated
non−targeted
PLK1
XIAP
E
Δ AUC MEKi vs p53 status
Wilcoxon, p = 0.001
TP53 status
mut
WT
Δ AUC MEKi vs RAS status
Wilcoxon, p = 0.17
RAS status

**Figure S3: Screening for synergistic effects between drugs and radiation, related to Figure 2. A,** Distribution of ΔAUCs of inhibitors stratified by mechanism of action. **B**, Distribution of ΔAUCs of MEKi, EGFRi and PARPi in individual organoid lines. **C**, Heatmap of ΔAUCs from the FDA-cancer library screen with ten rectal cancer organoids. **D,** Mean ΔAUCs vs. non-radiated AUCs according to TP53 and RAS status. **E,** Association of RAS and TP53 mutation status on MEKi ΔAUCs.

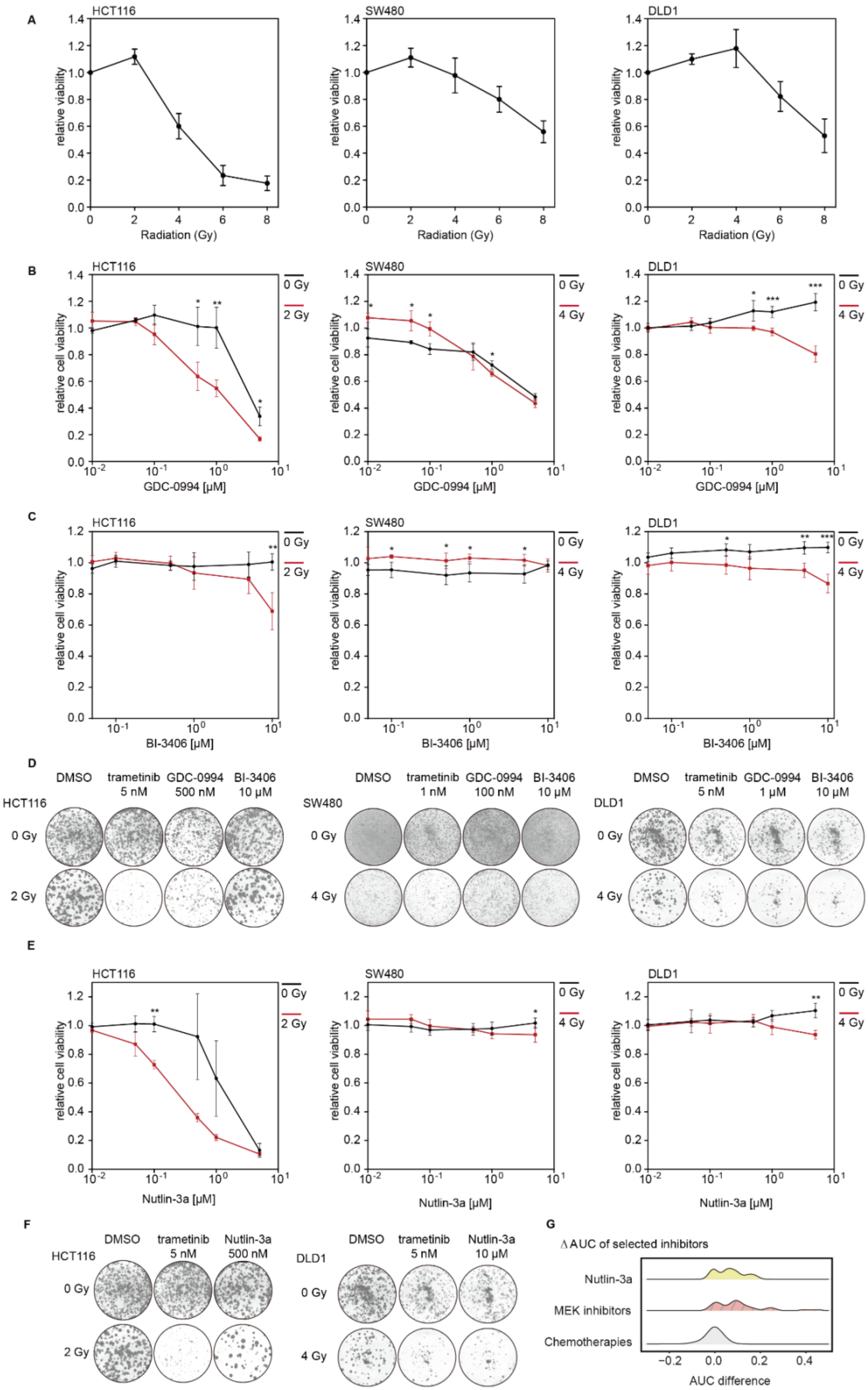
A
HCT116
SW480
DLD1
relative viability
Radiation (Gy)
B
relative cell viability
GDC-0994 [μM]
0 Gy
2 Gy
4 Gy
C
BI-3406 [μM]
D
DMSO
trametinib 5 nM
GDC-0994 500 nM
BI-3406 10 μM
trametinib 1 nM
GDC-0994 100 nM
GDC-0994 1 μM
E
Nutlin-3a [μM]
F
Nutlin-3a 500 nM
Nutlin-3a 10 μM
G
Δ AUC of selected inhibitors
Nutlin-3a
MEK inhibitors
Chemotherapies
AUC difference

**Figure S4: Effect of KRAS:SOS1 and ERK1/2 inhibitors on radiosensitivity of CRC cell lines, related to Figure 3. A**, Differential intrinsic radiosensitivity of CRC cell lines. **B-C**, Viability assay of CRC cell lines treated with increasing concentrations of ERK inhibitor GDC-0994 (B) or KRAS:SOS1 inhibitor BI-3406 (C) with and without radiation. Cell viability was determined after 5-6 days treatment by CellTiter-Glo. **D,** Colony forming assay of CRC cell lines treated with trametinib, GDC-009 or BI-3406 -/+ radiation. Scans of complete wells of standard six-well plates are shown (9.6 cm² per well). **E**, Viability assay of CRC cell lines treated with increasing concentrations of MDM2 inhibitor Nutlin-3a with and without radiation. Cell viability was determined after 5-6 days of treatment by CellTiter-Glo. **F**, Colony forming assay of CRC cell lines treated with Nutlin-3a -/+ radiation. Scans of complete wells of standard six-well plates are shown (9.6 cm² per well) **G**, Comparison of distribution of ΔAUC between Nutlin-3a, MEK inhibitors and chemotherapy in organoid drug-radiation screen.

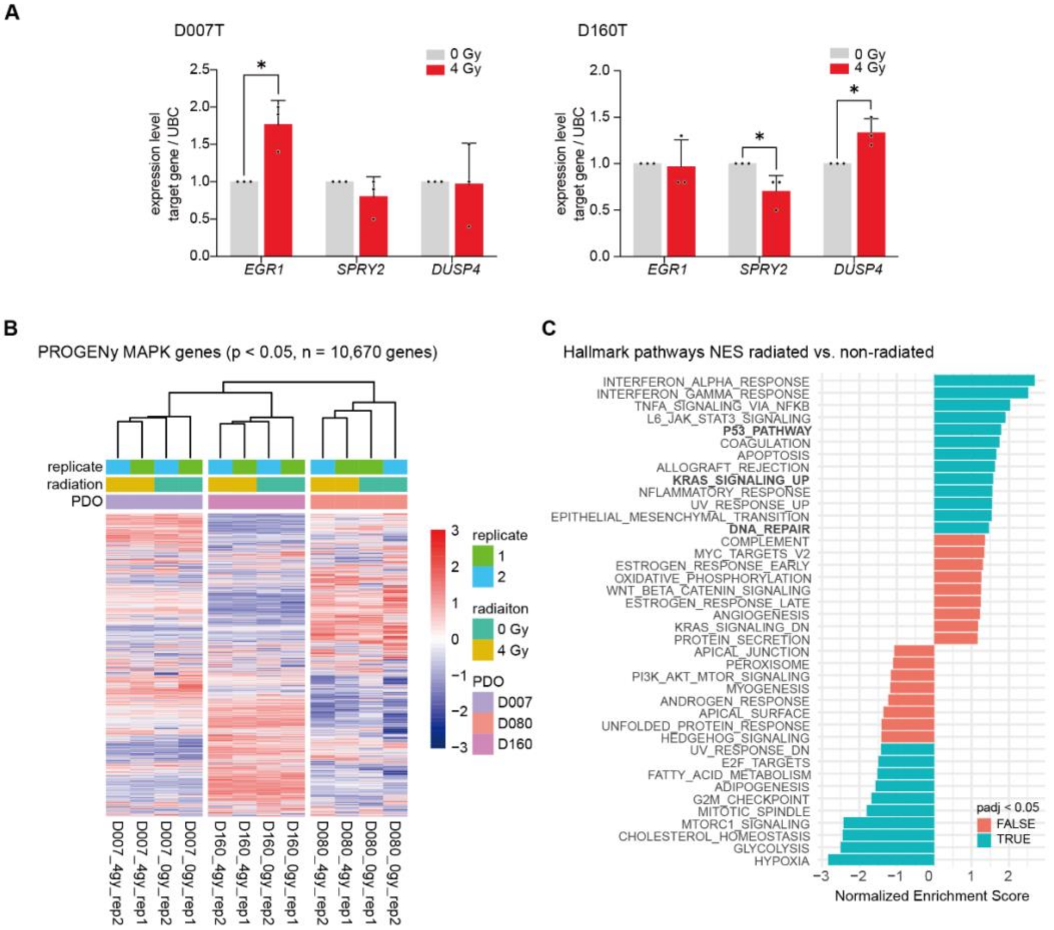


**Figure S5: Radiation/induced effects on signaling pathways, related to Figure 4.**

**A**, Transcriptional induction of target genes of RAS-MAPK pathway after radiation is suppressed by MEK inhibition in CRC organoids. Expression of genes is determined by qPCR. Data from three independent experiments are presented as mean ± SD. *p < 0.05, two-tailed Student's t-test. **B,** Unsupervised clustering of all 10,670 genes included in the PROGENy MAPK pathway gene signature list. Data clustering shows organoid line as the main driver of variance in the dataset. On the level of organoid lines, a clustering according to irradiated vs. non-irradiated condition can be observed. **C**, Gene set enrichment analysis of

HALLMARK gene sets in irradiated vs. non-irradiated organoids shows upregulation of gene sets “DNA_repair”, “P53_pathway” and “KRAS_signaling_up”, amongst others.

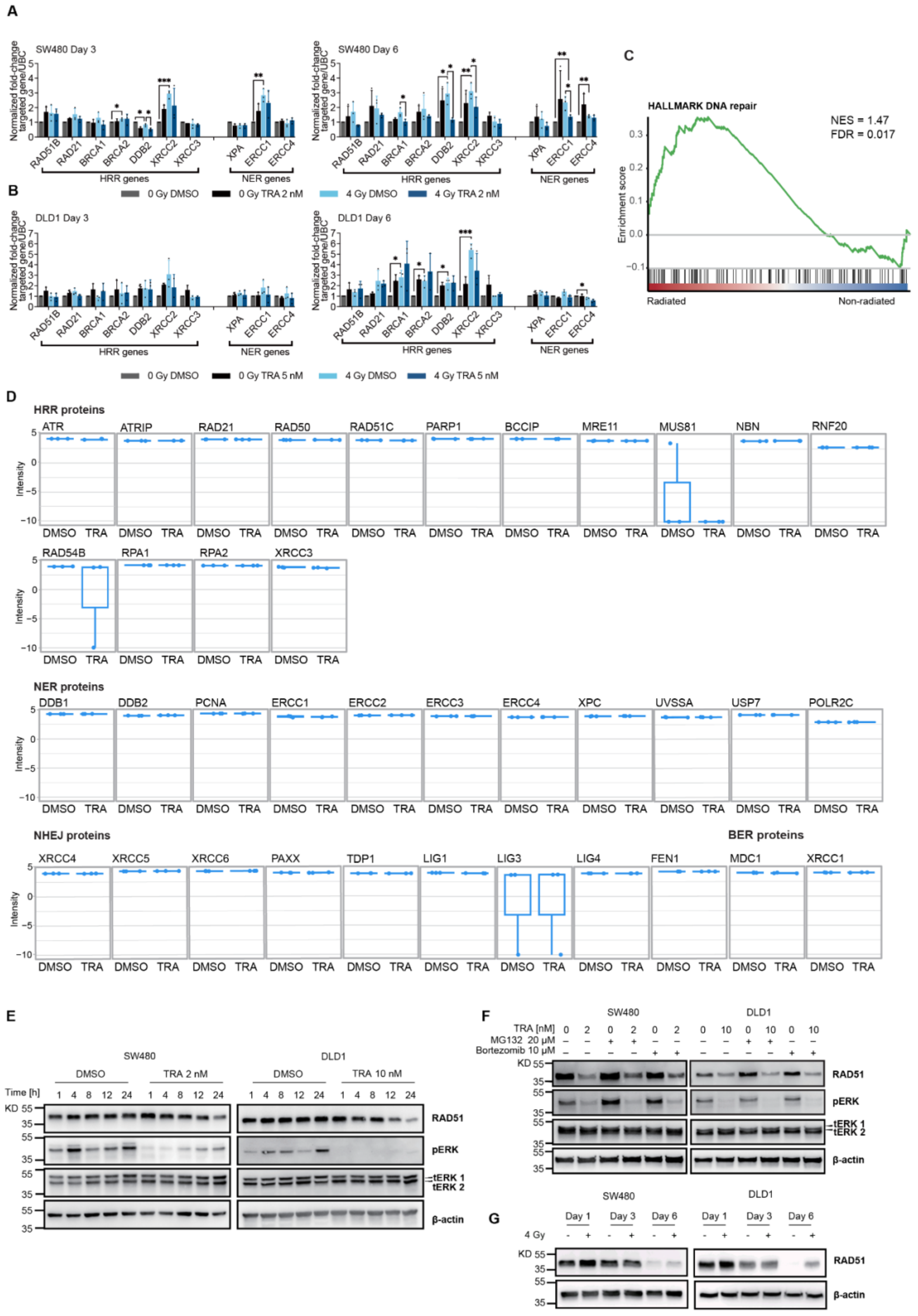
A
SW480 Day 3
SW480 Day 6
Normalized fold-change targeted gene/UBC
RAD51B
RAD21
BRCA1
BRCA2
DDB2
XRCC2
XRCC3
XPA
ERCC1
ERCC4
HRR genes
NER genes
0 Gy DMSO
0 Gy TRA 2 nM
4 Gy DMSO
4 Gy TRA 2 nM
B
DLD1 Day 3
DLD1 Day 6
0 Gy TRA 5 nM
4 Gy TRA 5 nM
C
HALLMARK DNA repair
NES = 1.47
FDR = 0.017
Enrichment score
Radiated
Non-radiated
D
HRR proteins
ATR
ATRIP
RAD21
RAD50
RAD51C
PARP1
BCCIP
MRE11
MUS81
NBN
RNF20
RAD54B
RPA1
RPA2
XRCC3
Intensity
DMSO
TRA
NER proteins
DDB1
DDB2
PCNA
ERCC1
ERCC2
ERCC3
ERCC4
XPC
UVSSA
USP7
POLR2C
NHEJ proteins
XRCC4
XRCC5
XRCC6
PAXX
TDP1
LIG1
LIG3
LIG4
FEN1
BER proteins
MDC1
XRCC1
E
SW480
DLD1
DMSO
TRA 2 nM
TRA 10 nM
Time [h]
KD
RAD51
pERK
tERK 1
tERK 2
β-actin
F
TRA [nM]
MG132 20 μM
Bortezomib 10 μM
G
Day 1
Day 3
Day 6
4 Gy

**Figure S6: Radiation and MEK inhibition-induced effects on DNA repair pathway in CRC, related to Figure 5.**

**A-B**, Radiation-induced transcriptional changes of homologous recombination repair (HRR) genes and nucleotide excision repair (NER) genes in CRC cell lines. Expression of genes is determined by qPCR. **C,** Barcode enrichment plot of HALLMARK gene set DNA_REPAIR in irradiated vs. non-radiated conditions. **D**, Protein levels of selected components of the DNA repair pathway (non-homologous end joining - NERJ, base excision repair - BER) in MEKi vs. DMSO treated HCT116 cells, determined by global proteomics analysis. Data from three independent replicates are shown. **E**, Time-dependent loss of RAD51 upon MEK inhibition occurs at 24 h after. **F,** Loss of RAD51 is not rescued by concomitant proteasome inhibition. **G**, Radiation increases RAD51 protein levels in CRC cell lines at late time points (day 6). **A-B,** Data from three independent experiments are presented as mean ± SD. $*p < 0.05$, $**p < 0.01$, $***p < 0.001$, two-tailed Student's t-test. **E-G**, Representative images of three independent biological replicates are shown.

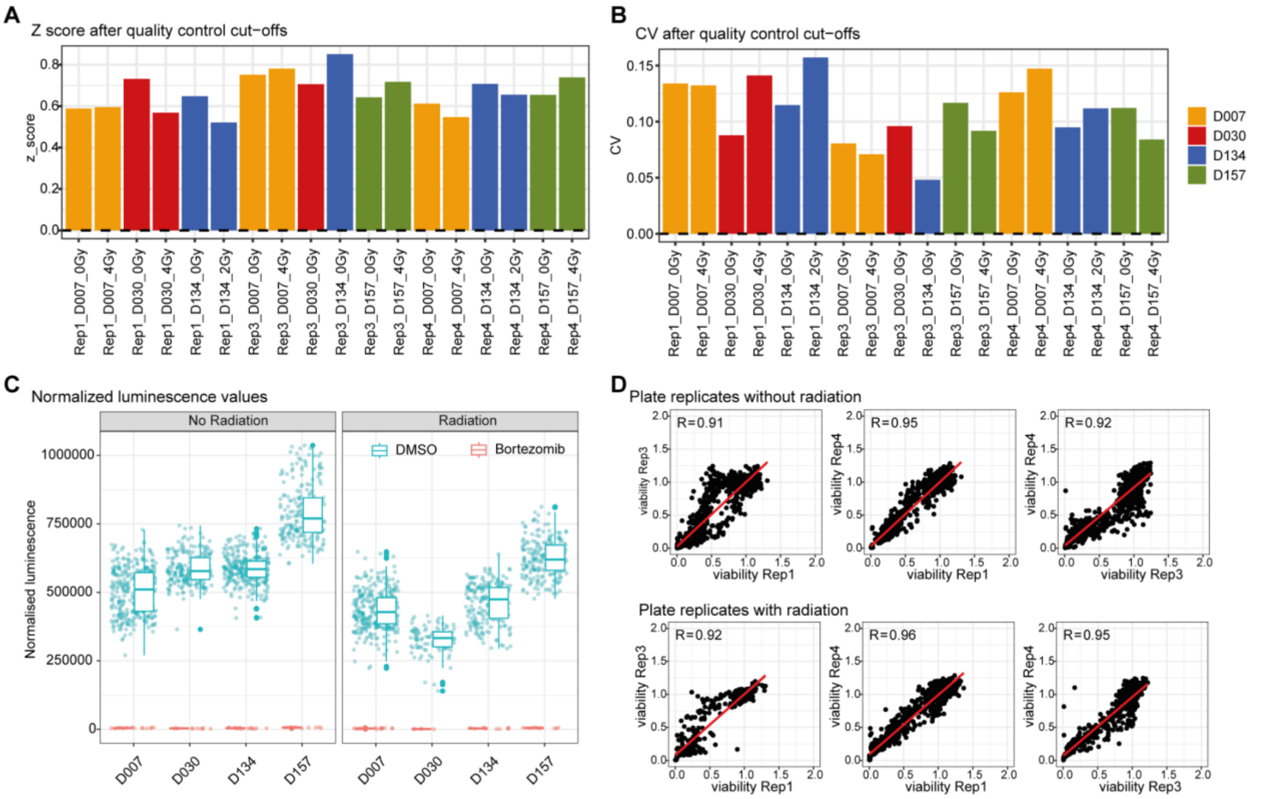


**Figure S7: PARP inhibitors synergize with MEK inhibitors in CRC models to enhance radiation response, related to Figures 6 - 7. A-D,** Quality controls of drug-drug-radiation profiling experiments. **A-B,** z-score and cv of all tested plates after quality control cut-offs were applied: two plates (D030T Rep. 3 4 Gy, D134T Rep 3 4 Gy) were excluded from further analysis. **C,** Normalized luminescence values of positive (bortezomib) and negative (DMSO) controls in radiation and non-radiation assays. **D,** Plate replicate correlations.

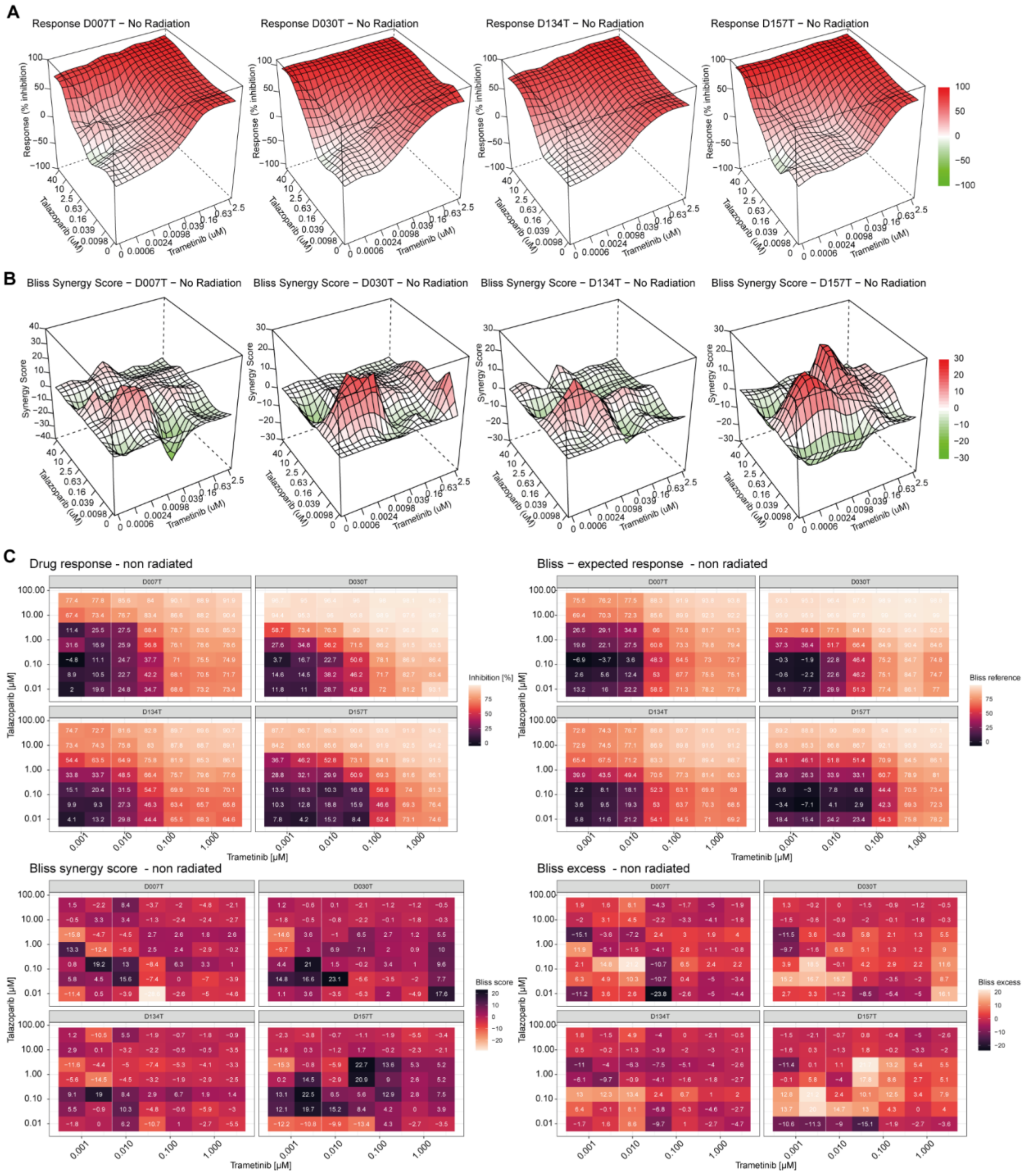


**Figure S8: PARP inhibitors synergize with MEK inhibitors in CRC models, related to Figure 6. A,** 3-dimensional response (% inhibition) surfaces of the talazoparib and trametinib combination in all four tested organoid lines. The surfaces contain fitted values. **D,** Bliss synergy surfaces of the talazoparib and trametinib combination in all four tested organoids. The surfaces contain fitted values. **C,** Drug response, Bliss expected response, Bliss synergy score and Bliss excess calculated for all dose combinations of talazoparib and trametinib in all four tested organoid lines. All data shown in this figure were obtained in absence of radiation treatment.

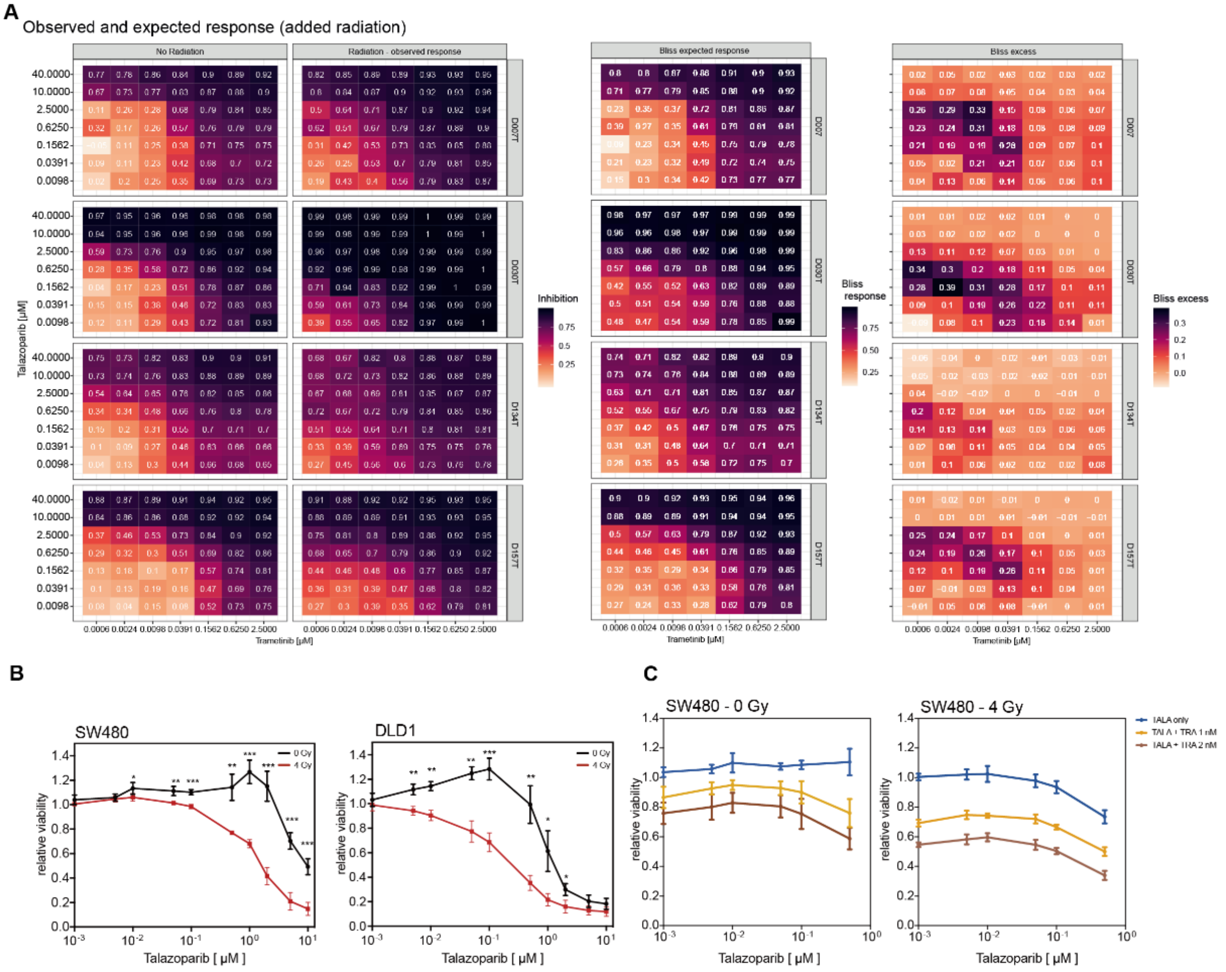


**Figure S9: PARP inhibitors synergize with MEK inhibitors to enhance radiation response, related to Figure 7.**

**A,** Response/inhibition matrix derived from talazoparib - trametinib combinations for all four tested organoid lines: Non-irradiated, irradiated, Bliss expected response, according to a model of added radiation to fixed combinations of trametinib and talazoparib, as well as Bliss excess (observed response - expected response) are shown. Data were normalized to non-irradiated DMSO controls. The complete matrices tested are shown for each drug. **B,** Viability assays of CRC cell lines treated with increasing concentrations of PARP inhibitor talazoparib with and without radiation. Cell viability was determined after 60 hrs of treatment by CellTiter-Glo. **C**, Viability assays of CRC cell line SW480 treated with increasing concentrations of PARP inhibitor talazoparib in combination with selected low-dose trametinib treatments, with and without radiation. Cell viability was determined after 60 hrs of treatment by CellTiter-Glo. **B-C**, Data from three independent experiments are presented as mean ± SD. *p < 0.05, **p < 0.01, ***p < 0.001, two-tailed t-test.
